# Primate-conserved Carbonic Anhydrase IV and murine-restricted Ly6c1 are new targets for crossing the blood-brain barrier

**DOI:** 10.1101/2023.01.12.523632

**Authors:** Timothy F. Shay, Erin E. Sullivan, Xiaozhe Ding, Xinhong Chen, Sripriya Ravindra Kumar, David Goertsen, David Brown, Jost Vielmetter, Máté Borsos, Annie W. Lam, Viviana Gradinaru

**Author notes:** Co-corresponding author: T.F.S., Co-corresponding author: V.G. These authors contributed equally to this work (Timothy F. Shay, Erin E. Sullivan, Xiaozhe Ding).

## Abstract

The blood-brain barrier (BBB) presents a major challenge to delivering large molecules to study and treat the central nervous system (CNS). This is due in part to the scarcity of effective targets for BBB crossing, the identification of which is the crucial first step of drug development. Here, we leveraged a panel of adeno-associated viruses (AAVs) previously identified through directed evolution for improved BBB transport to reverse engineer protein targets for enhanced BBB crossing. We identify both murine-restricted Ly6c1 and primate-conserved carbonic anhydrase IV (Car4; CA4) as novel receptors for crossing the BBB. We demonstrate how these receptors can unlock new experimental and computational target-focused engineering strategies by creating the enhanced Ly6c1-binding vector AAV-PHP.eC and by applying AlphaFold2-enabled in silico methods to rank capsids against identified receptors and generate capsid-receptor binding models. Here, with Car4, we add a completely new receptor to the very short list currently available for crossing the BBB in humans and, with Ly6c1, we validate a pipeline for receptor-targeted engineering. The identification of Car4/CA4 and structural insights from computational modeling provide new paths toward human brain-penetrant chemicals (drugs) and biologicals (including gene delivery).

## INTRODUCTION

The blood-brain barrier (BBB) presents a fundamental bottleneck to the development of effective research tools and therapeutics for the central nervous system (CNS)*(1–3)*. This structure, comprised mainly of brain endothelial cells*(2, 4)* requires large molecules to be delivered via invasive intracranial injections, technically challenging focused ultrasound*(5)*, or receptor-mediated transcytosis*(6, 7)*. Rational design of BBB-crossing large molecules has long been hampered by our imperfect understanding of the mechanisms involved in transcytosis, with only a handful of targets, such as transferrin receptor*(8–11)*, validated for research and therapies*(7, 12– 14)*.

Directed evolution is a powerful method for generating biomolecules with enhanced fitness for desired properties despite an incomplete understanding of the underlying biological systems*(15)*. Importantly, the outcomes of directed evolution libraries could in turn be used to unlock new biology by probing the mechanism of action for molecules with evolved properties. We have decided to apply this paradigm of reverse-engineering directed evolution hits to the accumulating wealth of data resulting from selective pressure on adeno-associated virus (AAV) libraries for CNS enrichment after systemic administration*(16–25)*.

One such improved rodent BBB-crossing AAV capsid is PHP.eB*(26)*, which we previously identified by Cre-recombination-based AAV targeted evolution (CREATE) method*(17)* on the parent capsid AAV9*(27, 28)*. Following systemic injection in genetically divergent mouse strains, capsids such as PHP.eB can either show potent CNS tropism (as in C57BL/6J, FVB/NCrl, and DBA/2J) or akin to AAV9 (as in BALB/cJ)*(29–31)*. In non-human primates (NHPs), PHP.eB and others display AAV9-like CNS infectivity*(29, 32–34)*. As AAVs have become the vector of choice for human gene therapies, including for therapies of the CNS*(35–37)*, complications from directly applying mouse-evolved AAVs in new genetic backgrounds contributed to a shift toward performing AAV directed evolution in NHPs. In so doing, researchers hope to increase the likelihood of identifying AAVs whose enhanced tropism will translate to humans.

As both the pre-clinical validation and, increasingly, the generation of engineered capsids occurs in NHPs however*(34, 38, 39)*, the animals’ scarcity*(40, 41)* and costs slow the identification of engineered capsids while the risk of NHP-specific AAVs entering clinical trials remains. Nevertheless, examples are beginning to accumulate of capsids that can cross both rodent and NHP BBB*(34)* but, of utter importance, also of at least one capsid reported to cross the macaque but not the rodent BBB*(39)*. Collectively, this diverse set of capsids engineered over the past decade by many groups represents, through their yet unexplored mechanisms of action, an unprecedented opportunity to start unraveling new targets for crossing the BBB across strains and species.

As patients treated with any AAV are likely to develop neutralizing antibodies toward most future AAVs and preclude them from future AAV treatments*(35, 36)*, methods to efficiently identify targets of brain-enhanced AAVs are critically necessary. Thus, NHP-optimized and validated AAV capsids that might not cross the human BBB are a concerning possibility. Here, we demonstrate a path forward by identifying protein targets or BBB-crossing mechanisms that may directly translate to human drug development. Using brain-enhanced engineered AAVs previously identified by directed evolution selections in mice*(20, 22, 24, 34)*, we validate a pipeline to reverse engineer targets for potent BBB crossing. Focusing on engineered AAVs whose enhanced CNS infectivity upon systemic injection in mice is conserved across both C57BL/6J and BALB/cJ strains*(20, 22, 24)*, we identified using an in vitro screening model with in vivo validation two novel receptors for enhanced BBB crossing by engineered AAVs: murine-restricted Ly6c1 and primate-conserved carbonic anhydrase IV (Car4; CA4).

To demonstrate how a known target can unlock new engineering strategies, (1) we created an enhanced Ly6c1-binding AAV variant, AAV-PHP.eC, (2) we utilized a new method for Automated Pairwise Peptide-Receptor Analysis for Screening Engineered AAVs (APPRAISE-AAV for short) that uses AlphaFold2*(42)* to screen peptides against potential receptors in silico and (3) we generated AAV-receptor interaction models, including the first high-resolution binding model of PHP.eB with Ly6a. Our experimental and computational pipeline for learning how evolved AAVs enact their enhanced BBB crossing tropisms demonstrates target-driven capsid engineering with Ly6c1 and establishes in vivo that carbonic anhydrase IV is a receptor target for enhanced CNS transduction by AAVs with translational promise.

## RESULTS

### Identification of engineered AAVs that do not utilize Ly6a

Ly6a is the receptor responsible for the enhanced CNS tropism of PHP.B and PHP.eB in mice*(43– 45)*, and one strain-specific SNP disrupts its GPI-anchored membrane localization (Fig. 1a)*(45)*. Previously, we applied our M-CREATE directed evolution selection platform to a library of AAV9 variants (containing seven-amino-acid insertions at position 588 of capsid variable region VIII), identifying a family of engineered AAVs with diverse CNS tropisms and a shared sequence motif, typified by their founding member, AAV-PHP.B (Fig. 1b)*(22)*. The variants’ enhanced CNS potency is lacking in BALB/cJ mice, a phenomenon explained by their shared reliance on Ly6a for BBB crossing*(43–45)*. Recently, multiple engineered AAVs outside the PHP.B sequence family which retain their enhanced CNS tropism in BALB/cJ mice were identified*(20, 22, 24)*. Before attempting to de-orphanize these AAVs, we first sought to confirm their independence of Ly6a.

**Fig. 1.**
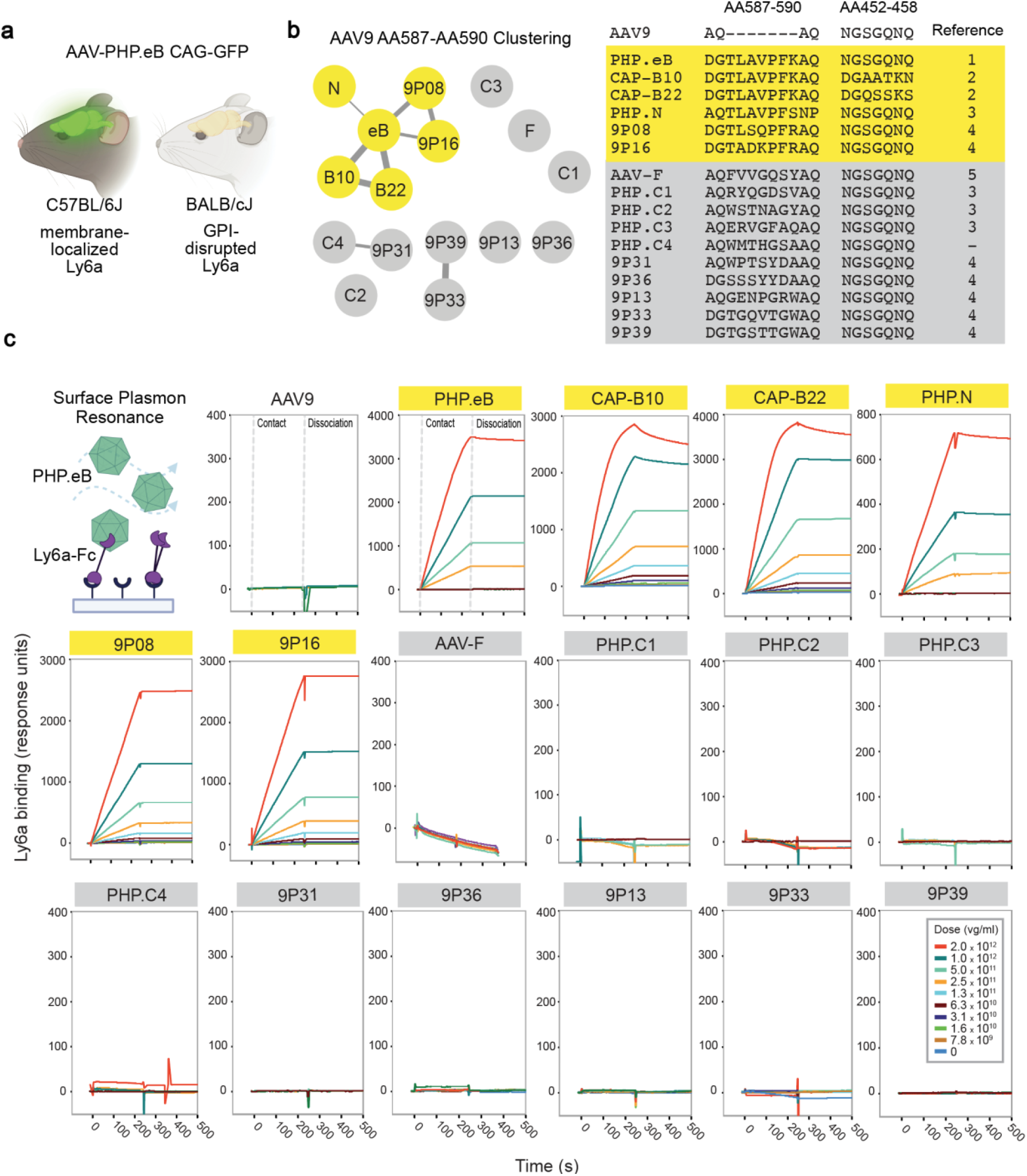
Identifying engineered AAVs that do not utilize Ly6a for BBB crossing. **a**, AAV-PHP.eB can efficiently cross the blood brain barrier using membrane-localized Ly6a in C57BL/6J mice but not GPI-disrupted Ly6a in BALB/cJ mice. **b**, Clustering analysis of CNS-tropic engineered AAV insertion sequences. Thickness of edges represents degree of relatedness between nodes. AAVs in yellow show a high degree of relatedness to PHP.eB. AAVs in gray show little relation to PHP.eB or each other. The AA sequences inserted between positions 588-589 and 452-458 of the AAV9 capsid for each variant is shown. References: 1: Chan et al. (2017), 2: Goertsen et al. (2022), 3: Ravindra Kumar et al. (2020), 4: Nonnenmacher et al. (2021), 5: Hanlon et al. (2019) **c**, Surface plasmon resonance (SPR) of engineered AAVs binding to Ly6a-Fc captured on a protein A chip. Data are representative of 2 independent experiments.

To directly probe potential Ly6a binding interactions, we performed surface plasmon resonance (SPR). To ensure the detection of even weak interactions, we dimerized Ly6a by fusion to Fc and immobilized it at high density on a protein A chip, and tested each AAV analyte across a range of concentrations (Fig. 1c). As expected, AAV9 showed no evidence of binding at any concentration, and PHP.B sequence family members all showed strong binding interactions with Ly6a. While precise affinities could not be determined due to the effects of avidity and mass transport, the interaction profiles were consistent with sub-nanomolar affinities. Conversely, all BALB/cJ-enhanced engineered AAVs were indistinguishable from AAV9, exhibiting no detectable interaction with Ly6a.

### Construction of a cell culture screen for putative receptors of engineered AAVs

Reasoning that a receptor that engineered AAVs can co-opt for efficient BBB crossing is likely to be both highly expressed and highly specific to the endothelial cells of the brain, we analyzed previously-collected single-cell RNA sequencing data from dissociated C57BL/6J brain tissue (Fig. 2a)*(46)*. We investigated gene expression levels within CNS endothelial cell clusters and differential expression compared to all other CNS cell type clusters. Genes were filtered to select only those annotated in Uniprot as localized to the plasma membrane before calculating their endothelial cell differential expression score in scanpy. This score was then plotted against each transcript’s mean abundance, revealing a long tail of highly expressed and highly specific CNS endothelial membrane proteins. Encouragingly, Ly6a appeared at the far reaches of this tail. From this analysis, we selected a panel of 40 abundant and specific candidate receptors (Table S1).

**Fig. 2.**
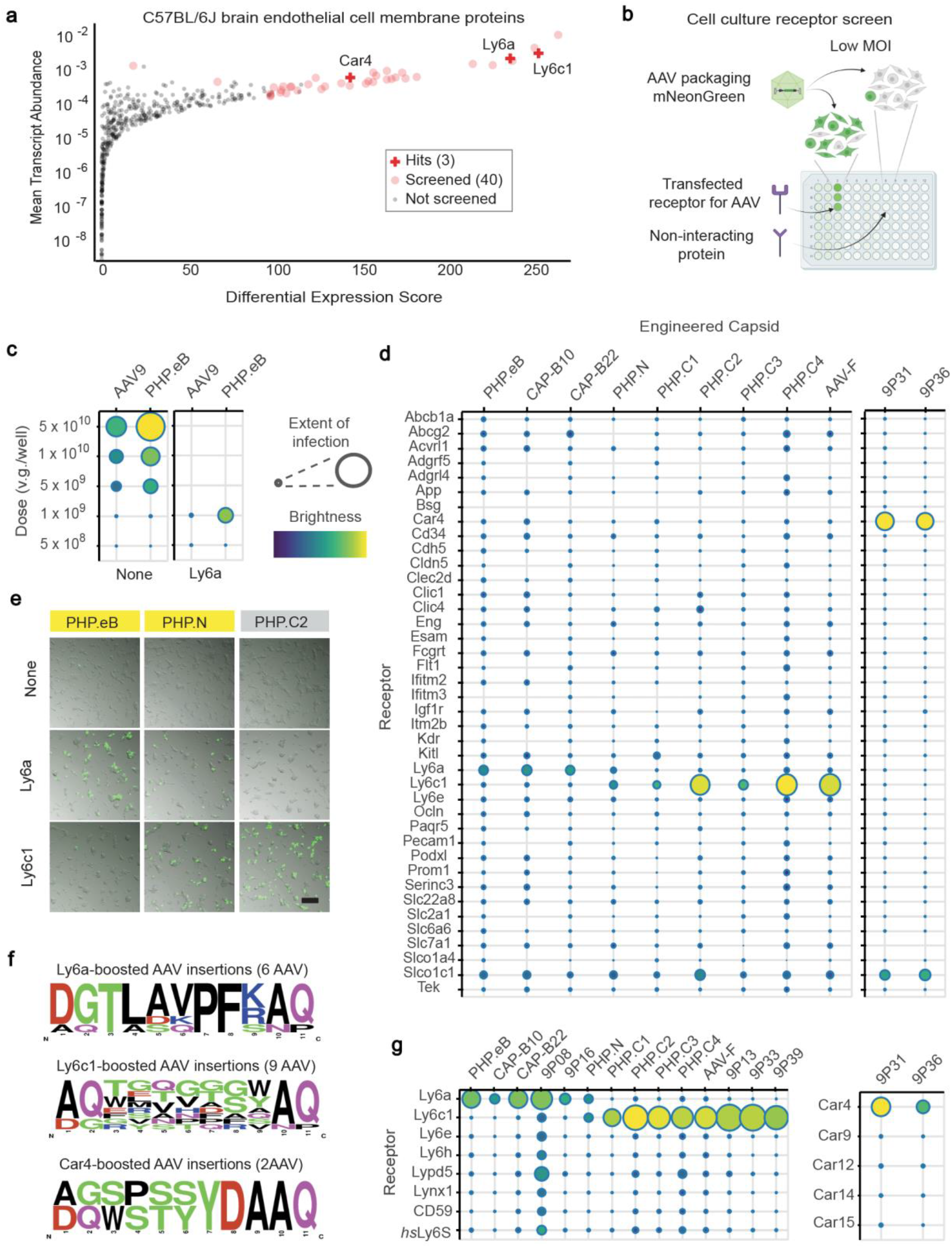
Novel receptors enhance infectivity of engineered AAVs *in vitro*. **a**, Plot of membrane protein differential expression score in endothelial cells calculated in scanpy versus mean transcript abundance in endothelial cells constructed from single-cell RNA sequencing of C57BL/6J cortex. Proteins selected for cell culture screening are indicated by red circles (see Supplementary Table 1 for full details) and hits are labeled and marked with red crosses. **b**, Potential receptors were screened in cell culture by comparing AAV fluorescent protein transgene levels at low multiplicity of infection (MOI) in mock-transfected cells and cells transfected with each potential receptor. **c**, Dose dependence of AAV9 and PHP.eB packaging CAG-mNeonGreen in HEK293T cells in 96-well plates. At 1 × 10^9^ v.g. per well, PHP.eB has markedly higher potency than AAV9 in Ly6a-transfected HEK293T cells. Scales show extent of infection (Max: 0.75, Min: 0.03) and total brightness per signal area (Max: 0.79, Min: 0.39) **d**, Potency of engineered AAVs for HEK293T cells transfected with the potential receptor panel. Extent of infection (*Left*, Max: 0.33, Min: 0.001; *Right*, Max: 0.55, Min: 0.03), Total brightness per signal area (*Left*, Max: 0.55, Min: 0.06; *Right*, Max: 0.61, Min: 0.11) **e**, Representative images of HEK cells cultured in 96 well plates, either mock-transfected (none) or transfected with a potential receptor, infected with 1 × 10^9^ v.g. per well of AAVs (yellow label background indicates Ly6a binding) packaging CAG-mNeonGreen and imaged 24 hours post transduction. An overlay of brightfield and fluorescence images is presented. Scale bar = 200 µm. **f**, Amino acid frequencies by position among engineered AAVs found to have infectivity enhancements with Ly6a (6 viruses), Ly6c1 (9 viruses), and Car4 (2 viruses). **g**, Potency of engineered AAVs for HEK293T cells transfected with Ly6 and Car family potential receptors. Extent of infection (*Left*, Max: 0.51, Min: 0.03; *Right*, Max: 0.52, Min: 0.05), Total brightness per signal area (*Left*, Max: 0.75, Min: 0.13; *Right*, Max: 0.79, Min: 0.34).

We and others have observed that expression of membrane-localized Ly6a in HEK293 cells selectively improves the potency of PHP.eB infection compared to AAV9 at low multiplicity of infection (MOI), with extent of infection and brightness of infected cells markedly increased (Fig. S1a)*(45)*. Hypothesizing that this property is likely to be conserved among BBB-crossing receptors, we made this behavior the basis of a receptor transient overexpression screen (Fig. 2b-c). We cloned the C57BL/6J coding sequence of each of the 40 candidate receptors into a mammalian expression plasmid and tested each against engineered AAVs in triplicate at two different doses (Fig. 2a, 2d). PHP.eB and Ly6a served as a positive control.

As expected, we found that all members of the PHP.B sequence family showed a marked boost in infectivity in Ly6a-transfected cells compared to untransfected cells, while Ly6a-independent AAVs performed identically in both conditions (Fig. 2d-e, & Fig. S1b). Interestingly, the infectivity of all Ly6a-dependent capsids was boosted to a similar extent (Fig. S2b).

### Identification of Ly6c1 and Car4 as receptors for BBB crossing of Ly6a-independent engineered AAVs

We observed boosts in infectivity for all tested Ly6a-independent AAVs with novel receptors (Fig. 2d). Surprisingly, given their different sequence families, all of the initial Ly6a-independent AAVs responded to the same candidate receptor, Ly6c1 (Fig. 2d-g and Fig. S1b). In addition, despite its Ly6a-dependent pattern of CNS infectivity across murine strains*(22)*, PHP.N also exhibited enhanced infectivity in Ly6c1-transfected cells. At a higher dose, PHP.N outperformed both AAV9 and PHP.eB in CBA/J mice (Fig. S3), which express a GPI-disrupted Ly6a. While polymorphisms in Ly6c1 exist between mouse strains (as shown by a published analysis of the 36 mouse strains for which whole genomes are available *(45)*), we found that, unlike Ly6a, none of the polymorphisms were predicted to disrupt GPI-anchoring of Ly6c1 to the plasma membrane using PredGPI*(47)*. To determine the specificity of the Ly6c1 interaction, we performed a follow-up screen with additional closely-related and CNS-expressed Ly6 superfamily members, and found no cross-reactivity of Ly6c1-dependent AAVs with other Ly6 proteins (Fig. 2g). CAP-B10 and CAP-B22, which have seven amino acid substitutions in capsid variable region IV and show enhanced potency in adult marmosets*(34)*, did not exhibit any additional receptor interactions that would explain their NHP tropism in either of the screens described above or a third screen with marmoset CNS-expressed Ly6 family members (Fig. S2c). Nor did any Ly6-interacting AAV cross-react with the recently described human Ly6S, a close relative of murine Ly6a (Fig. 2g)*(48)*.

We also expanded the Ly6 follow-up screen to include a second set of seven previously identified engineered AAVs (9P)*(24)*, which yielded additional five Ly6a and Ly6c1-interacting AAVs (Fig. 2g). The 9P AAVs 9P31 and 9P36, however, did not display enhanced infectivity with any Ly6 protein in the panel and so were tested on the full receptor screen (Fig. 2d). Both 9P31 and 9P36 displayed an infectivity boost with the GPI-linked enzyme carbonic anhydrase IV (Car4; CA4)(Fig. 2d and Fig. S1b). This interaction was specific to Car4 among membrane-associated carbonic anhydrases (Fig. 2g). As with Ly6c1, polymorphisms in Car4 across mouse strains are not predicted to impact GPI-anchoring to the plasma membrane*(45, 47)*. Unlike mouse-restricted Ly6a and Ly6c1, Car4 is conserved throughout vertebrates, including non-human primates and humans*(49–51)*. Therefore, we chose to confirm our screen results in vivo using Car4-knockout mice (B6.129S1-Car4tm1Sly/J, Jackson labs strain #008217). Immunofluorescence confirmed that Car4 is strongly expressed throughout the brain vasculature of homozygous WT/WT mice and completely absent in KO/KO (Fig. 3a). When dosed with 3 × 10^11^ v.g. per animal, PHP.eB, 9P31, and 9P36 all strongly expressed in both the brain and liver of wild-type mice (Fig. 3b). Under the same conditions in KO/KO mice, PHP.eB was unaffected whereas both 9P31 and 9P36 completely lack enhanced CNS tropism (Fig. 3c and Fig. S3b). Liver tropism, on the other hand, was decoupled from this effect, with all three viruses showing strong transduction in KO/KO mice. It is possible that any 9P31 and 9P36 potency differences in the periphery of KO/KO mice might be due to pharmacokinetic effects of no longer efficiently accessing the CNS.

**Fig. 3.**
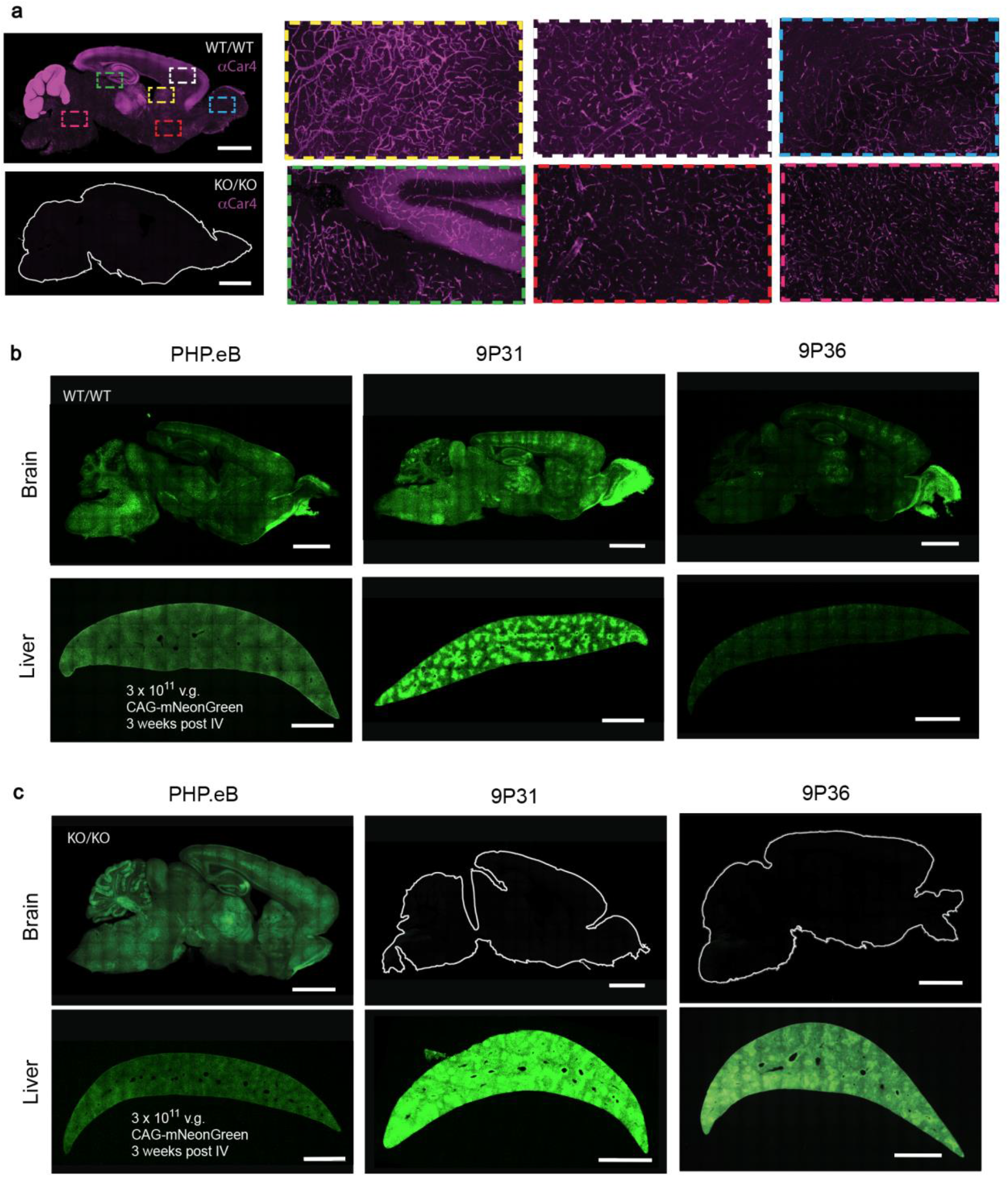
Carbonic anhydrase IV is required for the CNS potency of Car4-dependent AAV. **a**, Immunostaining for Car4 in the brains of WT/WT and KO/KO Car4 mice. Magnified regions from WT/WT demonstrate endothelial expression across diverse brain regions. **b**, AAV-PHP.eB, 9P31, and 9P36 packaging mNeonGreen under the control of the ubiquitous CAG promoter were intravenously administered to WT/WT Car4 mice at a dose of 3 × 10^11^ v.g. per animal (n=3 per condition). Three weeks after administration, transgene expression was assayed by mNeonGreen fluorescence throughout the brain and liver. **c**, AAV-PHP.eB, 9P31, and 9P36 packaging mNeonGreen under the control of the ubiquitous CAG promoter were intravenously administered to KO/KO Car4 mice at a dose of 3 × 10^11^ v.g. per animal (n=3 per condition). Three weeks after administration, transgene expression was assayed by mNeonGreen fluorescence throughout the brain and liver. Scale bars = 2 mm.

Of note, every AAV we tested showed a moderate boost in infectivity in cells transfected with Slco1c1 (also known as Oatp1c1), an integral membrane anionic transporter (Fig. 2d). Unlike for other potential receptors, however, the mNeonGreen signal was weak and diffuse, extending beyond cell boundaries (Fig. S1c), suggesting a transgene export or cell health phenotype. The universality of this effect and the specificity of Slco1c1 expression in the brain suggests a possible role in the weak BBB transcytosis of parent AAV9. None of the other 36 candidate receptors produced a meaningful infectivity boost for any of the 16 engineered AAVs screened.

### Directed evolution of an improved Ly6c1-dependent capsid

Next, we aimed to demonstrate a proof-of-concept for receptor-targeted directed evolution. For this purpose, we chose the murine receptor Ly6c1 (vs. the human Car4) due to prompt availability of (1) strains with clear BBB differences, e.g. Ly6a, and (2) diverse Cre-transgenic animals for M-CREATE selections that, for now, are not available for Car4 in other species. In addition, given the mixed backgrounds of preclinical animal models, it remains important to have mechanistically distinct gene delivery vectors for rodents. We therefore sought to engineer a single optimized capsid for broad adoption as a research tool in GPI-disrupted Ly6a mouse strains, a still unmet need, (Fig. 4) using our previously-developed M-CREATE method for AAV capsid directed evolution*(22)*. This method uses Cre-dependent AAV genome recovery from desired tissues and cell types after in vivo selection in Cre-transgenic mice and allows deep characterization of selected capsids across various cell types and tissues.

**Fig. 4.**
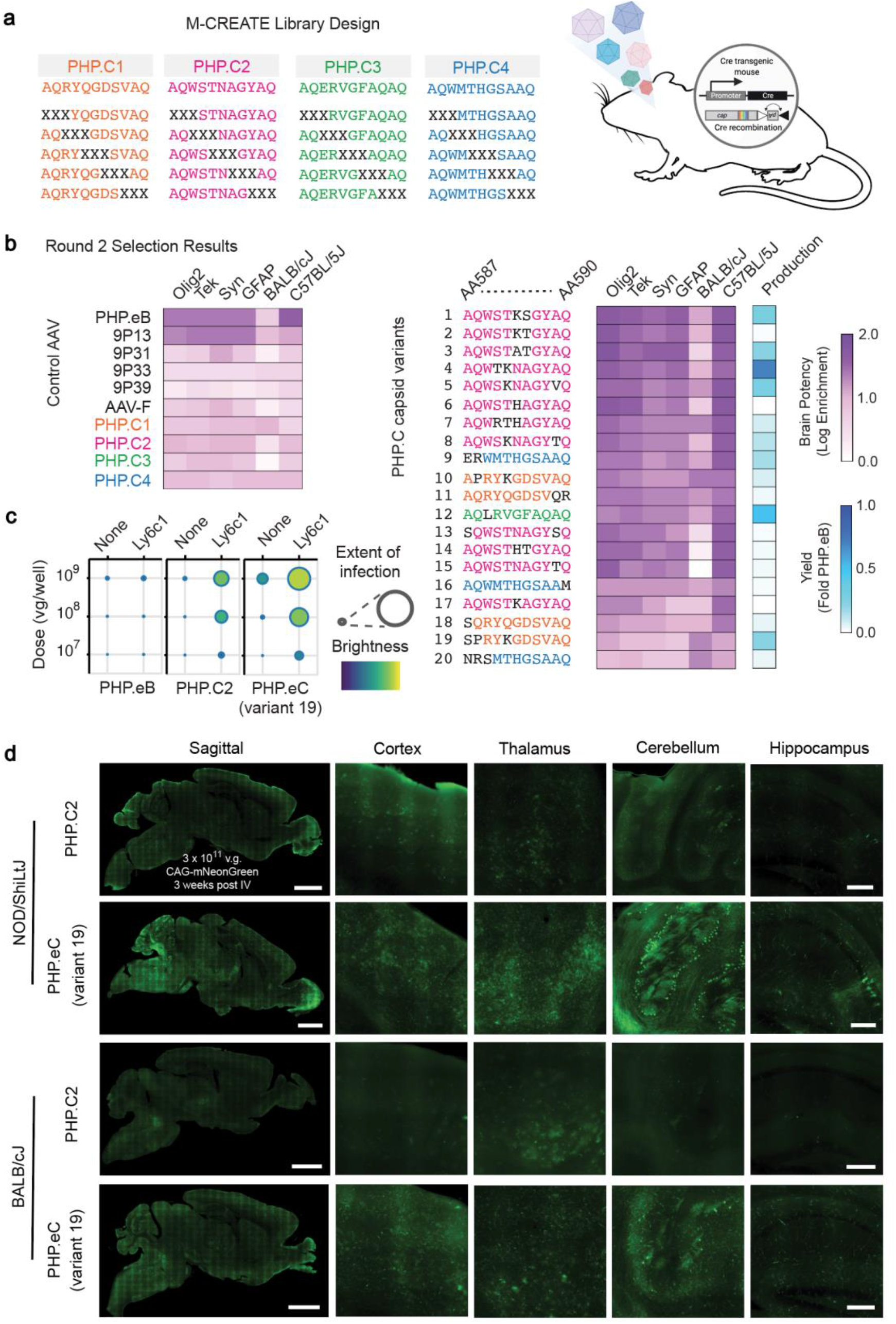
Engineering an enhanced Ly6c1-dependent AAV. **a**, AAV library design strategy. Scanning 3-mer libraries of PHP.C1 - C4 were constructed as shown with Xs indicating the position of NNK codons. AAVs were pooled and two rounds of M-CREATE selections were performed in 6-8 week old Cre-transgenic mice. **b**, Round 2 selection brain enrichments for diverse Cre-transgenic and wild-type mice of selected control AAVs and 20 capsid variants selected for further study. Yield was determined during small scale production for final screening across strains. **c**, Potency in cell culture infectivity assay of PHP.eB, PHP.C2, and PHP.eC (variant 19) in HEK293T cells transfected with mock (none) or Ly6c1 receptors, demonstrating that PHP.eC retains Ly6c1 interaction. Extent of infection (Max: 0.93, Min: 0.02), Total brightness per signal area (Max: 0.74, Min: 0.16) **d**, PHP.C2 and PHP.eC packaging mNeonGreen under the control of the ubiquitous CAG promoter were intravenously administered to NOD/ShiLtJ and BALB/cJ mice at a dose of 3 × 10^11^ v.g. per animal (n=3 per condition). Three weeks after administration, transgene expression was assayed by mNeonGreen fluorescence throughout the brain and liver, demonstrating PHP.eC’s increased potency. Sagittal image scale bars = 2 mm. Brain region scale bars = 250 µm.

Using M-CREATE, we constructed scanning 3-mer substitution capsid libraries in the chemically-diverse Ly6c1-interacting variants PHP.C1, PHP.C2, PHP.C3, and PHP.C4, and pooled these libraries for two rounds of selection in Syn-Cre mice (Fig. 4a). In the second round of selection, we also included Olig2-Cre, Tek-Cre, and GFAP-Cre mice, as well as wild-type C57BL/6J and BALB/cJ mice so that we could detect potential differences in enhancement between these strains or cell types during selection among the variants (Fig. 4b). Interestingly, while PHP.C2 variants dominated both rounds of selection in the C57BL/6J background (Cre-dependent or not), PHP.C1 variants dominated the round 2 Cre-independent selection in BALB/cJ (Fig. S4). Following selection, we individually produced and characterized top-performing variants. AAV-PHP.eC (variant 19), evolved from PHP.C1, retained Ly6c1 interaction in cell culture (Fig. 4c) and outperformed PHP.C2 in multiple mouse strains with membrane-disrupted Ly6a (Fig. 4d). PHP.eC thus provides a potent tool for transgene delivery in mouse strains without membrane-localized Ly6a.

### AlphaFold2-based methods to identify receptor-binding peptides and engineered AAV binding poses in silico

Having identified a panel of receptor and AAV capsid pairings, we aimed to see if we could capitalize on rapid advances in protein structure prediction to generate binding poses for engineered AAVs and their newly identified receptors. We began by applying an AlphaFold2-based computational method*(42, 52)* for Automated Pairwise Peptide Receptor Analysis for Screening Engineered AAV (APPRAISE-AAV for short). Inspired by recent work*(53, 54)*, this method uses AlphaFold2 to place surface-exposed peptides spanning mutagenic insertions (AA587 – 594) from two distinct AAV variants in competition to interact with a potential receptor (Fig. 5a). This comprises the minimal peptide to encompass the solvent-exposed residues of capsid variable region VIII. A combination of physical and geometric scoring parameters that include interface energy, binding angle, and binding pocket depth calculations are used to generate a peptide competition metric. Results from these individual pairwise competitions can be assembled into larger matrices that rank sets of AAV capsid insertion peptides according to their receptor-binding probability encoded in the AlphaFold2 neural network. When applied to Ly6a and our newly identified receptors, we found that the experimentally-verified Ly6a, Ly6c1, and Car4 insertion peptides rise to the top of their respective rankings (Fig. 5b-c). Some false negatives were also observed, however, as in 9P08 with Ly6a or 9P36 with Car4.

**Figure 5.**
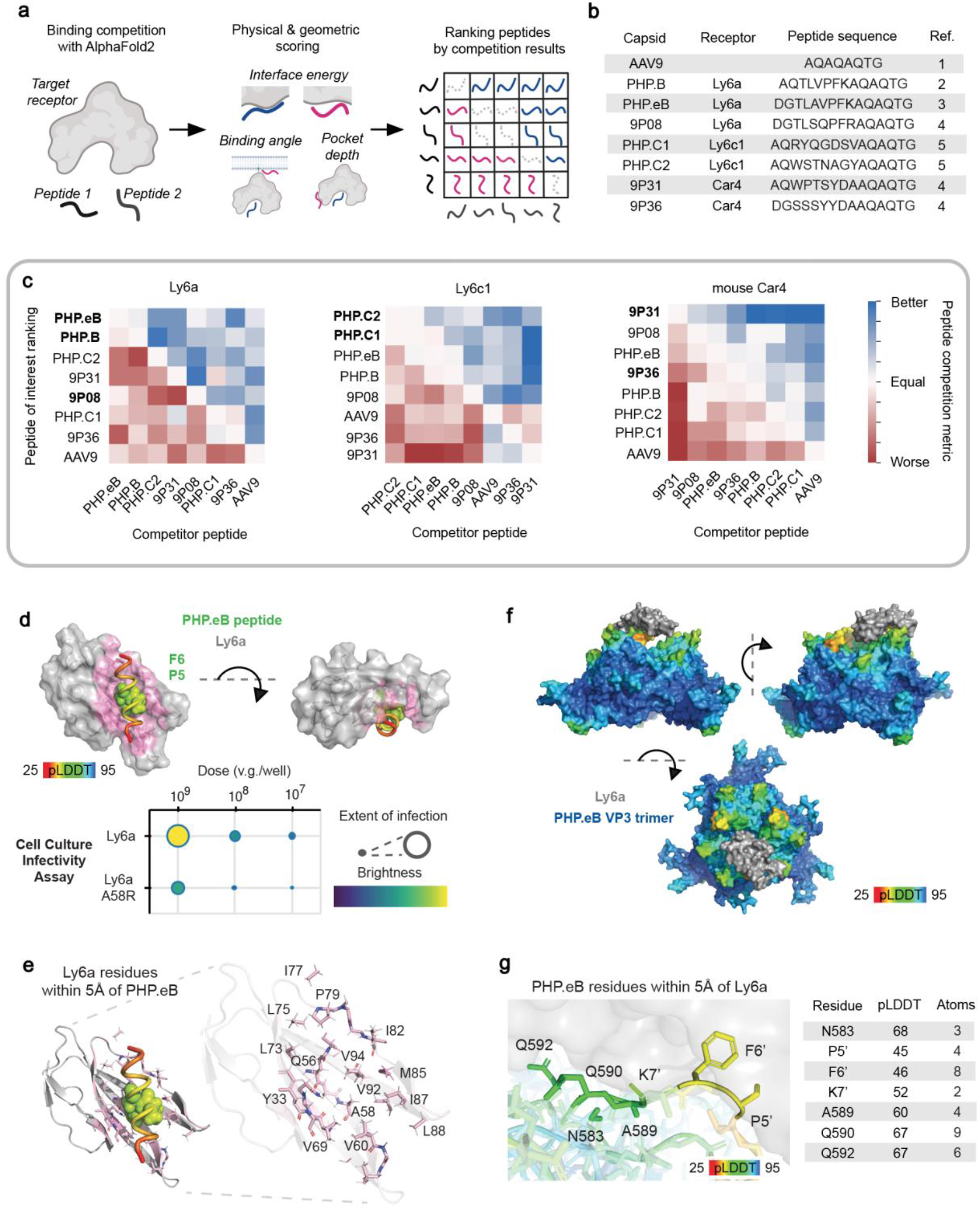
In silico predictions of engineered AAV receptors and Ly6a interaction complex pose. **a**, Overview of AlphaFold2-based in silico Automated Pairwise Peptide-Receptor Analysis for Screening Engineered AAVs (APPRAISE-AAV for short). Surface peptides from AAV variants are put in pairwise binding competition using AlphaFold2. A peptide competition metric is calculated according to each peptide’s interface energy, binding angle, and pocket depth (see Materials and Methods section for details) before being assembled into broader ranked matrices of interaction likelihood. Competition results reflect the relative peptide binding probability encoded in the AlphaFold2 neural network. **b**, Table of engineered AAV capsids, their confirmed receptor, and the capsid peptide sequence used in APPRAISE-AAV. References 1: Gao et al (2004), 2: Deverman et al (2016), 3: Chan et al (2017), 4: Nonnenmacher et al (2021), 5: Ravindra Kumar et al (2020) **c**, Matrices ranking AAV peptides by their average competition metric over ten replicate conditions for Ly6a, Ly6c1, and mouse Car4. AAV peptide labels in bold indicate those experimentally identified to interact with the corresponding receptor. Metric values out of range (−100 - 100) were capped to range limits. **d**, AlphaFold2-predicted Ly6a-PHP.eB peptide complex structure. PHP.eB peptide is colored by pLDDT (predicted Local Distance Difference Test) score, a per-residue estimate of the model confidence. The highest confidence side chains, P5’ and F6’, are shown as spheres. Ly6a A58R mutation, chosen to disrupt the predicted peptide interaction, resulted in reduced potency in the cell culture infectivity assay. Extent of infection (Max: 0.29, Min: 0.03), Total brightness per signal area (Max: 0.61, Min: 0.16). **e**, Ly6a residues with at least 2 atoms within 5 angstroms (Å) of the modeled PHP.eB peptide. **f**, Complete model of the PHP.eB trimer and Ly6a complex. The AlphaFold2 structural prediction from **d** was combined with a capsid monomer-receptor structural prediction and optimized using Rosetta Remodel within the context of the AAV trimer (Supplementary Figure 6a). **g**, Zoom-in view of the PHP.eB-Ly6a binding interface in modeled PHP.eB-Ly6a complex and PHP.eB residues with at least 2 atoms within 5 Å of Ly6a.

In addition to predictions of whether a peptide binds to a receptor, we can also computationally interrogate the structural details of the binding interaction. We generated binding poses by pairing the top AAV insertion peptide with its receptor and validated the binding pose for each pairing by repeating our cell culture screen with receptors containing point mutations hypothesized to disrupt the high-confidence region of the binding interface (as determined by the per-residue estimated model confidence pLDDT score and consistency between replicate models) in these predicted poses.

We began by modeling the interaction of PHP.eB with Ly6a so that the wealth of existing experimental data on this interaction*(55–57)* could be used to build confidence in our methods prior to their application in our newly discovered receptor interactions. The PHP.eB peptide is predicted to nestle in a groove in Ly6a, forming strong interactions at Pro5’ and Phe6’ (Fig. 5d) with several Ly6a residues (Fig. 5e). We therefore introduced a point mutation in this groove, Ly6a Ala58Arg, and found that it disrupts PHP.eB’s enhanced infectivity with the wild-type receptor. This experimental result further bolsters confidence in in silico APPRAISE-AAV rankings.

To gain a full picture of the AAV-receptor interaction, we next modeled the PHP.eB insertion peptide and Ly6a receptor complex within the context of the AAV capsid three-fold symmetry spike. This structure is challenging for standard modeling tools because of the large size of an AAV capsid (∼200kDa per trimer) as well as the often weak and dynamic binding interactions between engineered capsids and receptors (µM affinities possible without avidity*(56)*). AlphaFold2 failed to capture direct contact between full-length PHP.eB capsid and Ly6a in either a monomer-receptor or trimer-receptor configuration. To address this challenge, we developed an integrative structure modeling pipeline. In this pipeline, an initial model of an AAV capsid trimer predicted using AlphaFold2-Multimer*(52)* is structurally aligned with an AlphaFold2-predicted peptide-receptor complex model through the high-confidence Pro5’ and Phe6’ residues of the peptide insertion and RosettaRemodel*(58)* optimization of the linking peptide residues within the context of the AAV capsid three-fold symmetry spike (Fig. S5a). This complete binding model (Fig. 5e), provides a snapshot for a dynamic interaction that has thus far proven resistant to high-resolution structural characterization*(56)*.

The PHP.eB-Ly6a model coheres with available experimental results. RMSD between our PHP.eB monomer model and a cryo-EM-based model*(56)* (PDB ID: 7WQO) is 0.36 angstrom. RMSD increases in PHP.eB’s engineered loop to 1.36 angstroms. The only high-confidence deviation from cryo-EM structures of un-complexed PHP.eB is the side chain of Phe6’, which shows no significant electron density, indicating flexibility, but forms a stable interaction with Ly6a in our model (Fig. S5b, right). The high confidence prediction of Pro5’ and Phe6’ aligns with recent evidence showing that PFK 3-mer insertion alone is sufficient to gain Ly6a binding*(57)*. While Ly6a can bind any insertion loop of a trimer, additional interactions induce steric clashes supporting a ratio of one Ly6a per capsid trimer. Interestingly, a PHP.eB-Ly6a complex ensemble image forced to contain 60 bound copies of Ly6a resembles a recently reported CryoEM map, whose analysis pipeline would average over all 60 singly-occupied binding sites to form a composite map (Fig. S5c)*(56)*. Our model shows that a single copy of both Ly6a and AAVR PKD2 domain may bind to the same three-fold spike simultaneously without clashing (Fig. S5d), in agreement with saturation binding experiments*(56)*. Consistent with previous work showing the Ly6a SNP D63G does not affect PHP.eB binding*(45)*, the residue is greater than 10 angstroms from the PHP.eB peptide atoms in our models. The PHP.eB-Ly6a complex model includes several interactions involving AAV insertion-adjacent residues, which is consistent with a previous report (Figure 5g)*(55)*.

Having validated our methods against experimental data for the PHP.eB Ly6a interaction, we next applied these structural modeling methods to our newly identified receptors. Interestingly, unlike for Ly6a and Car4, the predicted binding pose for PHP.C2 peptide with Ly6c1 was found to vary with the version of AlphaFold-Multimer used, with v1 predictions closely matching mutational data from our cell culture infectivity assay (Fig. S5e). Such complementarity between versions has been reported previously*(59)*. In mouse Car4, 9P31 peptide invades the catalytic pocket of the enzyme (Fig. 6a). The 9P31 tyrosine residue shared with 9P36 approaches the enzyme active site and 9P31’s divergent tryptophan finds purchase in an ancillary pocket (Fig. 6b). This predicted binding pose is competitive with the binding site of brinzolamide (PDB ID 3NZC)*(60)*, a broad carbonic anhydrase inhibitor that is prescribed for glaucoma*(61)*. In our cell culture infectivity assay, brinzolamide shows a dose-dependent inhibition of 9P31 and 9P36 potency while PHP.eB is unaffected (Fig. 6b). The smaller brinzolamide binds deep in the catalytic core of Car4 where side chains are largely conserved between species (Fig. 6c). 9P31 peptide however extends to the surface of the enzyme where there is considerable sequence divergence that prevents cross-reactivity across species. Thus, while brinzolamide binds to both mouse and human Car4*(60, 62)*, 9P31 and 9P36 are selective for mouse Car4 (Fig. 6d). Chimeric receptors that swap a highly divergent loop of the 9P31 binding site show that this region is necessary but not sufficient to control 9P31 and 9P36 potency. Engineering a human CA4-binding AAV with optimal BBB crossing properties is both critically important and not trivial without ready *in vivo* model systems for validation, as illustrated by the extensive, multi-year efforts realizing transferrin receptor’s potential*(8, 9, 63–65)*.

**Fig. 6.**
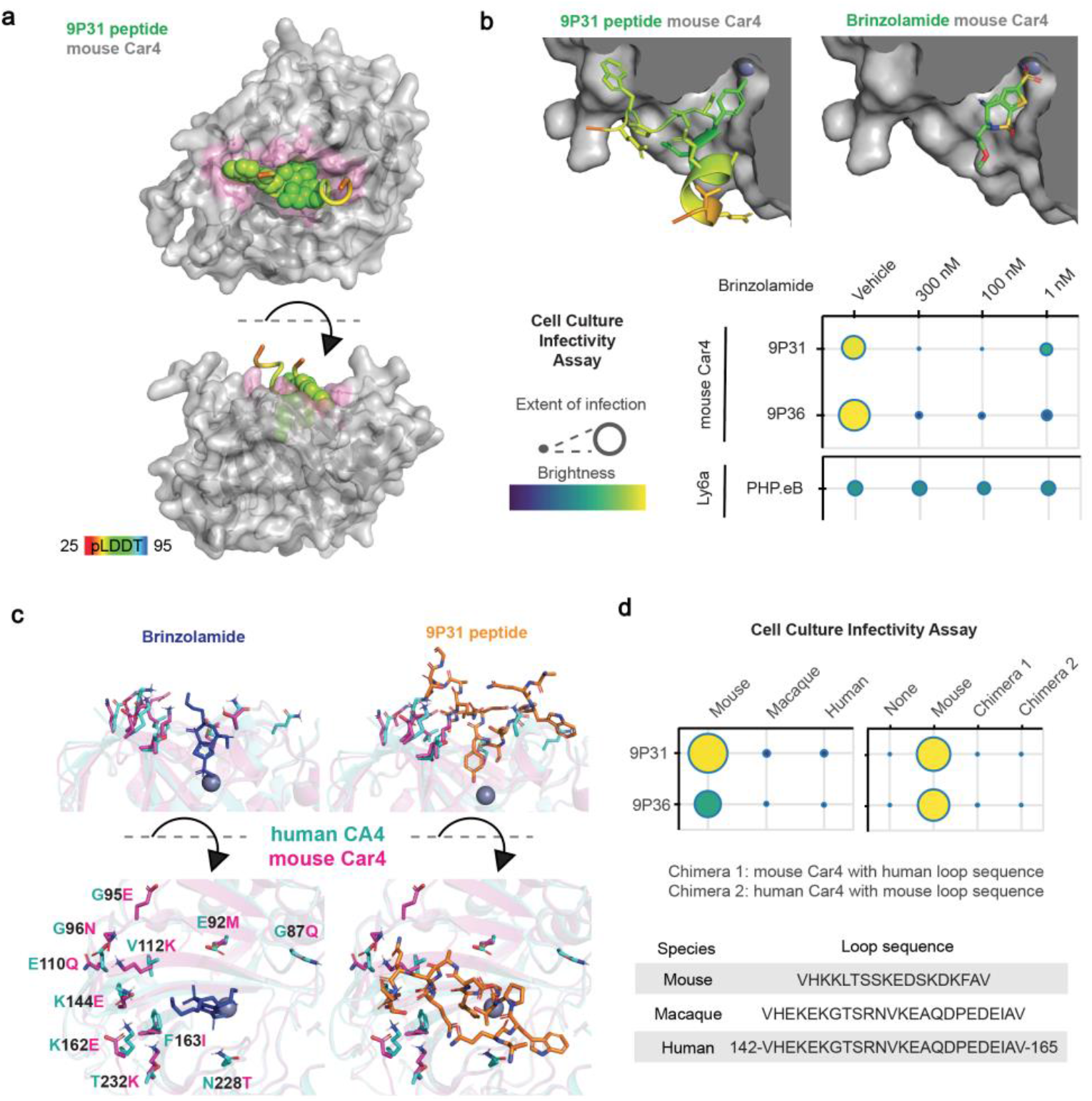
Engineered AAV interactions with carbonic anhydrase IV. **a**, AlphaFold2-predicted mouse Car4-9P31 peptide complex structure. 9P31 peptide is colored by pLDDT score at each residue with the highest confidence side chains shown as spheres. **b**, Cut-away view of mouse Car4 catalytic pocket with modeled 9P31 peptide binding pose (*top left)* and crystallographic brinzolamide binding pose (PDB ID: 3ZNC, *top right*). Cell culture infectivity assay of brinzolamide’s effects on engineered AAVs (*bottom*). Extent of infection (Max: 0.63, Min: 0.04), Total brightness per signal area (Max: 0.75, Min: 0.18) **c**, Views of amino acid side chains that differ between mouse (PDB ID: 3ZNC) and human (PDB ID: 1ZNC) carbonic anhydrase IV in relation to brinzolamide and 9P31 peptide binding poses. **d**, Potency in cell culture infectivity assay of 9P31 and 9P36 in HEK293T cells transfected with mouse, rhesus macaque, or human carbonic anhydrase IV receptors, as well as two chimeric receptors of mouse and human carbonic anhydrase IV that exchange the loop sequences depicted. Extent of infection (*left*, Max: 0.52, Min: 0.05, *right*, Max: 0.65, Min: 0.03), Total brightness per signal area (*left*, Max: 0.78, Min: 0.46, *right*, Max: 0.75, Min: 0.13)

## DISCUSSION

The blood-brain barrier restricts access to the CNS by large molecule research tools and therapeutics, limiting our ability to study and treat the brain*(1–4)*. Here we sought to expand the roster of protein targets through which biologicals and chemicals may access the CNS by de-orphanizing engineered AAVs selected through directed evolution for enhanced brain potency. While directed evolution methods have identified several engineered AAVs with enhanced tissue potency after systemic injection*(25)*, the mechanisms by which engineered AAVs gain their enhancements are, with a few recent notable exceptions*(38, 66)*, largely unknown. This is particularly true for engineered AAVs with enhanced potency in the CNS, where PHP.eB, which was found to use Ly6a in many mouse strains*(43–45)*, stands alone in being de-orphanized. The strain dependence and murine restriction of PHP.eB’s Ly6a interaction accelerated a push toward NHPs for engineered capsid identification and validation for translational vectors. However, human gene therapy’s increasing embrace of engineered AAV capsids in human clinical trials*(36, 37)* coupled with the scarcity and costs of NHP*(40, 41)*, highlight the need for higher through-put methods to validate novel AAVs with diverse, and conserved, mechanisms for crossing the BBB. By screening a curated pool of 40 candidate receptors selected for the intersection of their CNS expression level and endothelial-cell specificity, we were able to identify Ly6c1 and carbonic anhydrase IV as molecular receptors for enhanced blood brain barrier crossing of ten Ly6a-independent engineered AAVs (as well as Ly6a-dependent PHP.N). These findings allow for more efficient allocation of NHPs, inform future directed evolution library designs, and enable receptor-guided engineering directly for human protein interaction.

Interestingly, neither Ly6c1 nor Car4 had been identified as among the most enriched proteins in CNS endothelial cells compared to peripheral endothelial cells*(67)*. Given the distinct capsid sites for peptide insertion and galactose*(68)* or AAVR interaction*(69, 70)* and our model predicting simultaneous AAVR PKD2 and Ly6a interaction, it is likely that the receptors identified here work in concert with AAV9’s endogenous interaction partners to shape each AAV’s tropism.

While differences in their capsid insertion sequences suggested potentially diverse mechanisms for crossing the BBB, nine capsids were found to interact with the same receptor, Ly6c1. Differences in potency between AAVs in mouse strains can be dose-dependent, as we saw with PHP.N. This suggests that in some cases a cell culture screen can be sensitive to interactions that might only become functionally relevant *in vivo* at higher doses. Unlike Ly6a, none of the sequence polymorphisms are predicted to interfere with Ly6c1 GPI anchoring and thus membrane localization. Ly6c1 expression levels are also consistently high across in-bred mouse strains but are significantly lower in recently wild-derived mouse strains*(71)*. Together these features suggest that Ly6c1-utilizing AAVs may be useful research tools across genetically diverse mouse strains.

While we confirmed Ly6a interaction for CAP.B10 and CAP.B22, no additional interactions were identified. Thus, the mechanism by which these capsids (in contrast to PHP.B/eB) endow enhanced CNS potency in marmoset remains unclear*(34)*. It is possible that selecting a variable region IV library in the context of the PHP.eB variable region VIII insertion enriched for new receptor interactions outside the most abundant endothelial cell proteins and the Ly6 family. If a novel receptor can cooperatively enhance BBB crossing with Ly6a, it will enjoy outsized impact on CNS potency compared to acting alone. Thus, selection strategies promoting cooperativity may be employed to uncover lower likelihood receptors in mice and promote mechanistic diversity.

That two members of the Ly6 protein superfamily predominate as binding partners in CNS selections with AAV9 variable region VIII insertion libraries suggests a special complementarity between this library design and the Ly6 protein fold. This notion is supported by recent work suggesting additional interaction sites between wild-type AAV9 regions of AAV-PHP.eB and Ly6a*(55)*. Using our integrative modeling pipeline, we generated a complete, experimentally-validated receptor complex model for PHP.eB with Ly6a, which has otherwise resisted high-resolution structural characterization*(56)*. This model illustrates the complementarity of PHP.eB to Ly6a and predicts additional interactions outside of the insertion peptide. This insight provides opportunities for improved capsid engineering by both rational design (via *in vitro* selection for Ly6 family members with desirable expression patterns or conservation across species) and directed evolution (via negative selection pre-screens against purified Ly6 family proteins to encourage other BBB-crossing solutions). Our new APPRAISE-AAV *in silico* method is well suited to such screens. This method is also readily applied to any existing engineered capsid library dataset to mine for capsid variants likely to interact with a chosen target receptor, including Car4. Our modeling pipeline also provides high-confidence binding models for AAV receptor complexes that have proven difficult to structurally resolve. We note that the APPRAISE methodology is not limited to AAVs, and the pipeline for generating full AAV trimer complex structures may readily be employed to guide the translation of engineered peptide insertions identified through directed evolution in AAVs to other protein modalities.

In addition to the Ly6c1-interacting AAVs, our cell culture screen also identified two AAVs, 9P31 and 9P36, with pronounced infectivity boosts with carbonic anhydrase IV. Car4, also a GPI anchored protein, is known to localize to the luminal surface of brain endothelial cells throughout the cortex and cerebellum where it enzymatically modulates carbon dioxide-bicarbonate balance*(72, 73)*. While no specific role in BBB crossing has previously been attributed to Car4, it was recently found to be among the mouse proteins most strongly positively correlated with plasma-protein uptake in the brain (slightly stronger than the often-targeted transferrin receptor)*(74)*. This property was hypothesized at the time to be potentially useful for identifying receptors for enhanced BBB crossing*(75)*. Car4 is also expressed in the GI tract, kidney, and lung*(76, 77)*, as well as taste receptor cells where it allows us to sense carbonation*(78)*. While Ly6a expression in the kidney, heart, and liver*(79)* does not result in increased transduction there by PHP.B*(17)*, detailed peripheral characterization of 9P31, 9P36, and other Car4-interacting AAVs will be required to determine potentially enhanced infectivity in Car4-expressing peripheral tissue.

Carbonic anhydrase IV is broadly conserved across vertebrates and has similar CNS expression profiles in humans*(49–51)*, with recent single cell analyses of human brain vasculature confirming CA4’s expression in the human BBB*(80, 81)*. Thus, CA4-interacting AAVs are attractive candidates for translation across diverse model organisms and potentially in human gene therapies. A limitation of the current study, however, is that both 9P31 and 9P36 AAVs display enhanced potency with mouse Car4 but not rhesus macaque or human CA4. While neither virus would be expected to translate from mice to these species, we have identified a therapeutic target and mechanism for BBB crossing that may. The potential for specific engineered AAV binding epitopes to experience genetic drift between even closely related species confronts all products of directed evolution whose intended final use differs from their selection conditions. This potential takes on increasing importance when considering the potential for failed trials to preclude patients from future AAV treatments by eliciting cross-reactive neutralizing antibodies*(82–84)*. Future rational engineering of new AAVs against species-appropriate CA4, aided by our APPRAISE-AAV method, is a promising new avenue for the generation of non-invasive vectors with enhanced CNS potency. Targeting CA4 may also find application across diverse protein and chemical modalities.

In summary, using a convenient cell culture screen, we were able to identify diverse targets for enhanced BBB crossing by engineered AAV capsids. These include a novel candidate receptor for potent research tools in mice, Ly6c1, against which we evolved an enhanced capsid, AAV-PHP.eC, and a novel target that is broadly conserved across species, CA4, against which future engineered capsids may be designed for confident translation across species, including in humans. Cautioned by the differential performance of early engineered AAVs evolved in mice in hosts of different genetic backgrounds (with potentially different BBB compositions), the gene therapy field has increasingly transitioned to directed evolution in non-human primates to develop AAVs and other biologics. In addition to being significantly slower and more resource-intensive, the 25-30 million years of evolutionary divergence between macaques and humans*(85, 86)* may still prove a formidable barrier to human therapeutic translation, resulting in non-human primate capsids that might still fail in humans. Our studies highlight the importance of understanding engineered AAV mechanisms and demonstrate that high-throughput selections in rodents continue to play an important role in therapeutic target identification. By de-orphanizing mouse-selected AAVs, we identified carbonic anhydrase IV as a novel target for enhanced BBB receptor-mediated transcytosis across species, including humans.

## MATERIALS AND METHODS

### Plasmids

AAV capsid variants were subcloned into the pUCmini-iCAP-PHP.B backbone (Addgene ID: 103002). ssAAV genomes employed were pAAV:CAG-mNeonGreen (Addgene ID: 99134) and pAAV:CAG-2xNLS-EGFP (equivalent version with one NLS: Addgene ID: 104061), as noted in figures and legends. Receptor candidate plasmids were purchased from GenScript. All selected ORF clones were introduced to a pcDNA3.1+/C-(K)-DYK backbone, excepting Lynx1 in pcDNA3.1, which was generously shared by Prof. Julie Miwa (Lehigh). AAV capsid libraries were amplified from pCRII-9Cap-XE plasmid and subcloned into rAAV-ΔCap-in-cis-Lox2 plasmid for transfection with AAV2/9 REP-AAP-ΔCap (library plasmids available upon request from Caltech CLOVER Center)*(22)*.

### Animals

All animal procedures were approved by the California Institute of Technology Institutional Animal Care and Use Committee (IACUC) and comply with all relevant ethical regulations. C57BL/6J (000664), BALB/cJ (000651), CBA/J (000656), NOD/ShiLtJ (001976), Syn1-Cre (3966), GFAP-Cre (012886), Tek-Cre (8863), and Olig2-Cre (025567) mouse lines were purchased from Jackson Laboratory (JAX). Heterozygous Car4 knockout mice (008217) were cryo-recovered by JAX and bred at Caltech to generate homozygous WT/WT and KO/KO animals. Six to eight week old mice were intravenously injected with rAAV into the retro-orbital sinus. Mice were randomly assigned to a particular rAAV during testing of transduction phenotypes. Experimenters were not blinded for any of the experiments performed in this study.

### AAV vector production

AAV packaging and purification was performed as previously described*(31)*. Briefly, rAAV were produced by triple transfection of HEK293T cells (ATTC, CRL-3216) using polyethylenimine. Media was collected at 72 h and 120 h post-transfection and virus was precipitated in 40% polyethylene glycol in 2.5 M NaCl. This was resuspended and combined with 120 h post-transfection cell pellets at 37 °C in 500 mM NaCl, 40 mM Tris, 10 mM MgCl2, and 100 U mL-1 salt-active nuclease (ArcticZymes, 70910-202). The resulting lysate was extracted from an iodixanol (Cosmo Bio USA, OptiPrep, AXS-1114542) step gradient following ultracentrifugation. Purified virus was concentrated and buffer exchanged with phosphate buffered saline (PBS) prior to titer determination by quantitative PCR.

### AAV vector administration, tissue processing, and imaging

AAV vectors were administered intravenously to adult mice via retro-orbital injection at doses of 1 × 10^11^ or 3 × 10^11^ viral genomes (v.g.) as indicated in figures and legends. After three weeks of expression, mice were anesthetized with Euthasol (pentobarbital sodium and phenytoin sodium solution, Virbac AH) and transcardially perfused with roughly 50 mL of 0.1 M PBS, pH 7.4, and then another 50 mL of 4% paraformaldehyde (PFA) in 0.1 M PBS. Organs were then harvested and post-fixed in 4% PFA overnight at 4 °C before being washed and stored in 0.1M PBS and 0.05% sodium azide at 4 °C. Finally, the brain was cut into 100 um sections on a Leica VT1200 vibratome. Images were acquired with a Zeiss LSM 880 confocal microscope using a Plan-Apochromat x10 0.45 M27 (working distance, 2.0 mm) objective and processed in Zen Black 2.3 SP1 (Ziess) and ImageJ software.

### Single-cell RNA sequencing analyses

Analyses were performed on a pre-existing C57BL/6J cortex single-cell RNA sequencing dataset with custom-written scripts in Python 3.7.4 using a custom fork off of scVI v0.8.1, and scanpy v1.6.0 as described previously*(46)*. Briefly, droplets that passed quality control were classified as ‘neurons’ or ‘non-neurons’ using a trained scANVI cell type classifier, retaining only those cells above a false discovery rate threshold of 0.05 after correction for multiple comparisons. Non-neuronal cells were further subtyped using a trained scVI model and clustered based on the learned latent space using the Leiden algorithm as implemented in scanpy. Endothelial cell clusters were assigned if they were positive for all marker genes for that cell subtype. Membrane proteins were filtered by Uniprot keyword ‘cell membrane’ and differential expression scores were calculated in scanpy.

### Immunofluorescence

Immunofluorescence experiments were performed on HEK293T cells to label transiently transfected receptors such as Ly6a (Abcam ab51317, 1:200 dilution), Ly6c1 (Abcam ab15627, 1:200 dilution), and Carbonic Anydrase IV (Invitrogen PA5-47312, 1:40 dilution). HEK293T cells were seeded at 80% confluency in 6-well plates and maintained in Dulbecco’s Modified Eagle Medium (DMEM) supplemented with 5% fetal bovine serum (FBS), 1% non-essential amino acids (NEAA), and 100 U/mL penicillin-streptomycin at 37⁰C in 5% CO2. Membrane associated receptor candidates were transfected by polyethylenimine (PolySciences # 23966). Cells were seeded on Neuvitro Poly-D-lysine coated sterile German glass coverslips (Fisher Scientific #NC0343705) 24 hours post-transfection in 24-well plates then fixed in 4% paraformaldehyde once attached. Coverslips were blocked with 1X tris-buffered saline (TBS) containing 3% bovine serum albumin (BSA) for 30 minutes and incubated in primary antibody in 1x TBS, 3% BSA, and 0.05% Triton X-100 for 60 minutes at ambient temperature. Coverslips were washed three times in 1x TBS then incubated with secondary (Ly6a & Ly6c1: Invitrogen A-21247, 1:1000 dilution; Car4: Invitrogen A-21432, 1:1000 dilution) in the same medium for 60 minutes. Coverslips were mounted on slides with Diamond Antifade Mounting Media with DAPI (Invitrogen P36931). Fluorescent microscopic images were captured on a confocal laser-scanning microscope (LSM 880, Carl Zeiss, USA).

### Cell culture characterization of rAAV vectors

HEK293T cells were seeded at 80% confluency in 6-well plates and maintained in Dulbecco’s Modified Eagle Medium (DMEM) supplemented with 5% fetal bovine serum (FBS), 1% non-essential amino acids (NEAA), and 100 U mL^-1^ penicillin-streptomycin at 37 °C in 5% CO2. Membrane associated receptor candidates were transiently expressed in HEK293T cells by transfecting each well with 2.53 µg plasmid DNA. Receptor-expressing cells were transferred to 96-well plates at 20% confluency and maintained in FluoroBrite™ DMEM supplemented with 0.5% FBS, 1% NEAA, 100 U mL-1 penicillin-streptomycin, 1x GlutaMAX, and 15 µM HEPES at 37 °C in 5% CO2. Cells expressing each receptor candidate were transduced with engineered AAV variants at 1 × 10^9^ v.g. well^-1^ and 5 × 10^8^ v.g. well^-1^ in triplicate. Plates were imaged 24 hours post-transduction with the Keyence BZ-X700 using the 4x objective and NucBlue™ Live ReadyProbes™ Reagent (Hoechst 33342) to autofocus each well.

### Cell culture fluorescence image quantitation

All image processing was performed using our custom Python image processing pipeline, available at github.com/GradinaruLab/in_vitro_transduction_assay. In brief, the area of cells is determined in both brightfield and signal images and the percent of cells transduced and brightness per transduced area is determined from these images.

First, background subtraction is performed on the brightfield images by applying gaussian blur (skimage.filters.gaussian, sigma = 30, truncate = 0.35) and subtracting the product from the original brightfield image. In brightfield images, cells are silhouetted by the lamp producing both bright and dark edges. Histogram based thresholding can be applied to these images to determine bright and dark regions of the brightfield image, which can be combined to create a mask of cell edges in the image. Cells can be filled by applying skimage.morphology.closing, which runs a template over the image to fill contiguous regions (skimage.morphology.disk, radius = 2). The total area of cells in the brightfield image can then determined by summing all the pixels in the mask.

On the signal images, background subtraction is performed by applying gaussian blur (skimage.filters.gaussian, sigma = 100). Subtracting the product of gaussian blur from the original signal image produces an image with minimal fluctuations in background intensity. Histogram based thresholding is applied to this image to identify the intensity of background in the brightfield image and create a mask of bright regions in the image, which is comprised of transduced cells. Noise can be removed from the mask using skimage.morphology.remove_small_objects (min_size = 5). From this the total area of transduced cells can be determined by summing all the pixels in the mask.

After performing this segmentation, the percentage of cells transduced can be determined by taking the ratio of signal area to the total cell area. By multiplying the mask by the original image and summing all the pixel intensities in the product image, the total brightness of transduced cells can be determined. This value can then be divided by the total area of transduced cells to determine the brightness per transduced area.

### Receptor protein production

Ly6a-Fc was produced in Expi293F suspension cells grown in Expi293 Expression Medium (Thermo Fisher Scientific) in a 37 °C, 5% CO2 incubator with 130 rpm shaking. Transfection was performed with Expifectamine according to manufacturer’s instructions (Thermo Fisher Scientific). Following harvesting of cell conditioned media, 1 M Tris, pH 8.0 was added to a final concentration of 20 mM. Ni-NTA Agarose (QIAGEN) was added to ∼5% conditioned media volume. 1 M sterile PBS, pH 7.2 (GIBCO) was added to ∼3X conditioned media volume. The mixture was stirred overnight at 4 °C. Ni-NTA agarose beads were collected in a Buchner funnel and washed with ∼300 mL protein wash buffer (30 mM HEPES, pH 7.2, 150 mM NaCl, 20 mM imidazole). Beads were transferred to an Econo-Pak Chromatography column (Bio-Rad) and protein was eluted in 15 mL of elution buffer (30 mM HEPES, pH 7.2, 150 mM NaCl, 200 mM imidazole). Proteins were concentrated using Amicon Ultracel 10K filters (Millipore) and absorbance at 280 nm was measured using a Nanodrop 2000 spectrophotometer (Thermo Fisher Scientific) to determine protein concentration.

### Surface plasmon resonance

Surface plasmon resonance (SPR) was performed using a Sierra SPR-32 (Bruker). Ly6a-Fc fusion protein in HBS-P+ buffer (GE Healthcare) was immobilized to a protein A sensor chip at a capture level of approximately 1200-1500 response units (RUs). Two-fold dilutions of rAAVs beginning at 2 × 10^12^ v.g. mL^-1^ were injected at a flow rate of 10 µl min-1 with a contact time of 240 s and a dissociation time of 600 s. After each cycle, the protein A sensor chip was regenerated with 10 mM glycine pH 1.5. Kinetic data were double reference subtracted.

### Automated Pairwise Peptide Receptor Analysis for Screening Engineered AAVs (APPRAISE-AAV)

FASTA-format files containing a target receptor amino acid sequence (mature protein part only) as well as peptide sequences corresponding to amino acids 587 through 594 (wild-type AAV9 VP1 indices) from two AAV capsids of interest were used for structural prediction using a batch version of ColabFold*(87)* (alphafold-colabfold 2.1.14), a cloud-based implementation of multiple sequence alignment*(88–90)*, and AlphaFold2 Multimer*(52)*. The ColabFold Jupyter notebook was run on a Google Colaboratory session using GPU (NVIDIA Tesla V100 SXM2 16GB; we found that the same model of the GPU yielded the most consistent results). We chose alphafold2-multimer-v2 as the default AlphaFold version unless otherwise specified. Each model was recycled three times, and ten models were generated from each competition. Models were quantified with PyMol (version 2.3.3) using a custom script to count the total number of atoms in the interface (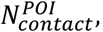, defined by a distance cutoff of 5 angstroms (Å)), the total number of atoms in the peptide that are clashing with the receptor (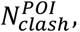, defined by a distance cutoff of 1 Å), the binding angle of the peptide (θ, defined as the angle between the vector from receptor gravity center to receptor anchor and the vector from receptor gravity center to peptide gravity center), and the binding depths (d, defined as the difference of the distance between the closest point on the peptide to the receptor center and the minor radius of the ellipsoid hull of the receptor normalized by the minor radius) of the peptide in each putative peptide-receptor complex model. The minor radii of the ellipsoid hulls of receptors were measured using HullRad 8.1 89 (Ly6a: 13.4 Å, Ly6c1: 12.7 Å, mouse Car4: 23.0 Å). Finally, the metric Δ*B*^*POI,competitor*^ for ranking the propensity of receptor binding was calculated by subtracting the total binding score of the peptide of interest by the counterpart score of the competing peptide:

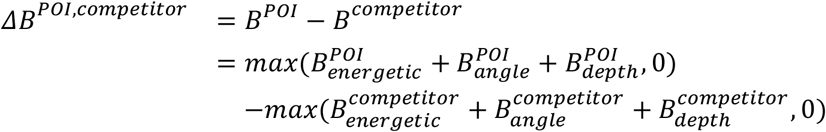

where the individual terms are defined as follows:

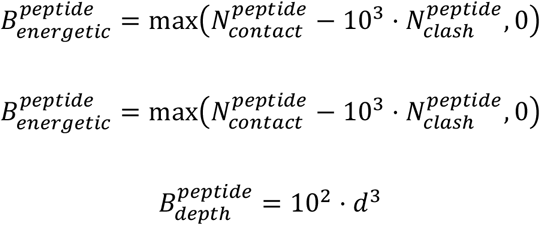

The mean number of this metric across replicates was used to form a matrix and plot a heatmap. Peptides in the heatmap were ranked by the total number of competitions each peptide won minus the total number of competitions it lost (competitions with Δ*B*^*POI,competitor*^) scores that have p-values greater than 0.05 in the one-sample Student’s t-test were excluded).

### Computational structure modeling of receptor-AAV complexes

Peptide-receptor structures were modeled using a similar procedure as described in the APPRAISE-AAV section but with only one single peptide of interest in the input file to achieve higher accuracy.

AAV trimer-receptor complex models were produced using an integrative structure modeling method (Fig. S5a). Trimers at the AAV three-fold symmetry interface were chosen as the minimal complete binding interface with a putative receptor that might recapitulate the entire viral particle while optimizing computational efficiency. First, a peptide-receptor model was generated by modeling the 15-mer peptide sequence between the residues 587 and 594 (both in wild-type AAV9 VP1 indices) from the AAV variant of interest in complex with the target receptor as described above. Then, a trimer model of the AAV variant of interest was modeled using AlphaFold2 Multimer. The two residues with the highest confidence score (pLDDT score) in the 15-mer peptide of the peptide-receptor model, Pro5’ and Phe6’, were structurally aligned to the corresponding residues on the first chain of the trimer model. A coarse combined model was then generated by combining the receptor and the two high-confidence AAV residues from the peptide receptor model with the remaining AAV residues from the trimer model. The two loops between Pro5’ and Phe6’ and the high-confidence AAV9 backbone in the coarse combined model (corresponding to residues 588-(588+)4’ and residues (588+)7’-590, respectively) were then individually remodeled using RosettaRemodel*(58)* from the Rosetta software bundle (release 2018.48.60516). Finally, these remodeled loops were merged to generate a final model. The pLDDT scores for each residue from the original AlphaFold2 outputs were used to color images of the final model.

### M-CREATE selections for PHP.eC

AAV capsid libraries were produced, administered, and recovered post-selection in vivo for next generation sequencing as described previously*(22)*. Briefly, an initial library was generated by pooling three amino acid NNK substitutions that scan from AAV9 587 through the 7mer insertion to AAV9 position 590 of AAV-PHP.C1, C2, C3, and C4 (Fig. 4a). A custom-designed synthetic round 2 library containing degenerate codon duplicates of 5515 capsid variants identified from the round 1 library post-selection and spike-in controls was synthesized in an equimolar ratio by Twist Biosciences. To prevent capsid mosaicism, only 10 ng of assembled library was transfected per 150 mm dish of 293T producer cells and assembled capsids were purified at 60 h post transfection. Round 1 library was retro-orbitally injected at 5 × 10^10^ v.g. per mouse in Syn-Cre mice, while round 2 library was retro-orbitally injected at 5 × 10^11^ v.g. per mouse in Syn-Cre, GFAP-Cre, Tek-Cre, Olig2-Cre, wild-type C57BL/6J, and wild-type BALB/cJ mice.

Two weeks post-injection, mice were euthanized and all organs including brain were collected, snap frozen on dry ice, and stored at −80 °C. Frozen tissue was homogenized using Beadbug homogenizers (Homogenizers, Benchmark Scientific, D1032-15, D1032-30, D1033-28) and rAAV genomes were Trizol extracted. Purified rAAV DNA (Zymo DNA Clean and Concentrator Kit D4033) was amplified by PCR using Cre-dependent primers, adding flow cell adaptors around the diversified region for next generation sequencing. NGS data was aligned and processed as described previously to extract round 1 variant sequence counts and round 2 variant enrichment scores.

## Acknowledgments

We thank Catherine Oikonomou for help with manuscript editing, Michael Altermatt for assisting in filtering scRNAseq datasets for membrane proteins, Min Jee Jang for designing RNA sequencing variant barcodes, Damien A. Wolfe for assisting in mouse perfusion and tissue collection, Josette Medicielo for plasmid purification, and Julie Miwa (Lehigh) for sharing a lynx1 expression plasmid. We thank the entire Gradinaru lab and CLOVER center staff for helpful discussion. Figures were created using images from BioRender.com. This work was primarily supported by NIH PIONEER DP1OD025535 (to V.G.) and the Beckman Institute for CLARITY, Optogenetics & Vector Engineering Research (CLOVER) for technology development and dissemination (to T.F.S. and V.G.).

## Author contributions

T.F.S. and V.G. conceived the project. T.F.S. and V.G. wrote the manuscript and prepared figures with input from all authors. X.D. and A.W.L. optimized Ly6a-Fc expression protocol and A.W.L. produced Ly6a-Fc protein. T.F.S. and E.E.S. produced AAVs. T.F.S. and J.V. performed SPR experiments. T.F.S. and D.B. analyzed the scRNAseq dataset. T.F.S. and E.E.S. developed and E.E.S. performed the cell culture infectivity assay. D.G. developed and implemented the in vitro transduction quantification and plotting pipeline, performed data analysis, and, with T.F.S., prepared in vitro transduction quantification plots. E.E.S. and M.B. performed immunofluorescence experiments. E.E.S., X.C., and M.B. performed Car4-KO experiments, X.D. performed APPRAISE-AAV and developed computational structural modeling strategies. T.F.S., S.R.K., E.E.S, and X.C. performed and analyzed AAV-PHP.eC selections and tested AAVs in wild type mice. T.F.S. and V.G. supervised and V.G. funded the project.

## Data and materials availability

The NGS datasets that are reported in this article will be made available with SRA accession codes listed here prior to final publication. Single-cell RNA sequencing datasets are available at CaltechDATA (data.caltech.edu/records/2090). Plasmids or viruses not available at Addgene may be requested from the Caltech CLOVER Center (clover.caltech.edu). The AAV-PHP.eC sequence will be made available with GenBank accession codes listed here prior to final publication. The code used for M-CREATE data analyses are available on GitHub: https://github.com/GradinaruLab/mCREATE. The code used for single-cell RNA sequencing analyses are available through GitHub on a different repository: https://github.com/GradinaruLab/aavomics. All in vitro transduction image processing was performed using our custom Python image processing pipeline, available at: https://github.com/GradinaruLab/in-vitro-transduction-assay.

