## Supplementary Material for "Primate-conserved Carbonic Anhydrase IV and murine-restricted Ly6c1 are new targets for crossing the blood-brain barrier"

### Supplementary Materials

| Name | Accession No. | Mean Expression | Differential Expression |
|---|---|---|---|
| Abcb1a | NM_011076.2 | 0.000688904 | 7.94237100 |
| Abcg2 | NM_011920.3 | 0.000745241 | 5.56071470 |
| Acvrl1 | NM_009612.3 | 0.000485079 | 6.78613330 |
| Adgrf5 | NM_001357332.1 | 0.000546496 | 7.45316400 |
| Adgrl4 | NM_133222.3 | 0.000763887 | 8.24139400 |
| App | NM_001198823.1 | 0.000845707 | 0.04600084 |
| Bsg | NM_009768.2 | 0.013027801 | 4.68415070 |
| Car4 | NM_007607.2 | 0.000713242 | 3.71078540 |
| Cd34 | NM_133654.3 | 0.000467251 | 3.09839130 |
| Cdh5 | NM_009868.4 | 0.000332645 | 7.48680500 |
| Cldn5 | NM_013805.4 | 0.005443705 | 9.17483200 |
| Clec2d | NM_053109.3 | 0.000584914 | 6.37040600 |
| Clic1 | NM_033444.2 | 0.000377934 | 2.64024160 |
| Clic4 | NM_013885.2 | 0.000408817 | 3.14403030 |
| Eng | NM_007932.2 | 0.000646813 | 5.99654530 |
| Esam | NM_027102.3 | 0.000660755 | 6.04634760 |
| Fcgrt | NM_010189.3 | 0.000503876 | 4.02808500 |
| Flt1 | NM_010228.3 | 0.002267079 | 8.53988600 |
| Ifitm2 | NM_030694.1 | 0.000314049 | 3.51603910 |
| Ifitm3 | NM_025378.2 | 0.001015794 | 5.31335400 |
| Igf1r | NM_010513.2 | 0.000750016 | 3.37413700 |
| Itm2b | NM_008410.2 | 0.001596154 | -0.10450481 |
| Kdr | NM_010612.2 | 0.000304256 | 6.83441160 |
| Kitl | NM_013598.3 | 0.000382663 | 3.28041740 |
| Ly6a | NM_010738.3 | 0.002695857 | 8.70076600 |
| Ly6c1 | NM_010741.3 | 0.003812270 | 8.84188900 |
| Ly6e | NM_008529.3 | 0.001050145 | 2.95476170 |
| Ocln | NM_008756.2 | 0.000208863 | 6.83609100 |
| Paqr5 | NM_028748.2 | 0.000262998 | 7.77428870 |
| Pecam1 | NM_008816.3 | 0.000423881 | 7.13435360 |
| Podxl | NM_013723.3 | 0.000407256 | 6.36354730 |
| Prom1 | NM_001163577.1 | 0.000271125 | 5.45687630 |
| Serinc3 | NM_012032.4 | 0.000562215 | 1.23178270 |
| Slc22a8 | NM_001164634.1 | 0.000243451 | 5.05416900 |
| Slc2a1 | NM_011400.3 | 0.001843255 | 5.43814370 |
| Slc6a6 | NM_009320.4 | 0.001148025 | 3.84664630 |
| Slc7a1 | NM_007513.4 | 0.000403344 | 4.17060850 |
| Slco1A4 | NM_030687.1 | 0.001701704 | 8.39613200 |
| Slco1c1 | NM_021471.2 | 0.000828427 | 3.68831500 |
| Tek | NM_013690.3 | 0.000224058 | 7.49632500 |

**Table S1: Single-cell RNA sequencing profiles of the C57BL/6J brain endothelial cell membrane proteins included in cell culture screen.**

### Supplementary Figures

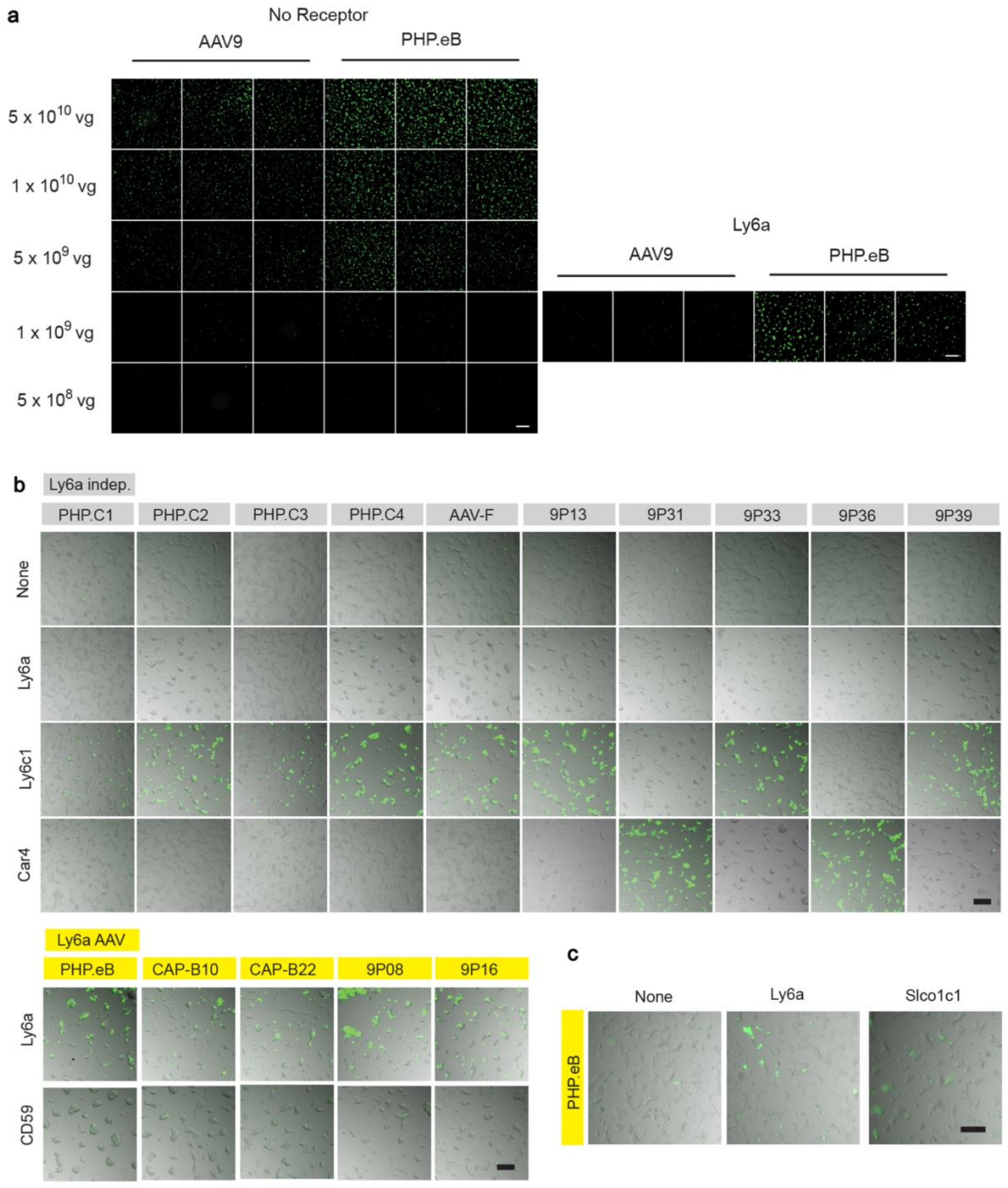


**Fig. S1. Representative images from cell culture infectivity assay. a,** Fluorescence images showing the dose dependence of AAV9 and PHP.eB packaging CAG-mNeonGreen in HEK293T cells in 96-well plates. At 1 x $10^9$ v.g. per well, PHP.eB has markedly higher potency than AAV9 in Ly6a-transfected HEK293T cells. Representative images are shown. Scale bars = 500 μm. **b,** Representative images of HEK cells cultured in 96-well plates, either mock-transfected (none) or transfected with a potential receptor, infected with 1 x $10^9$ v.g. per well of PHP.B sequence family AAVs (yellow label background indicating Ly6a binding) or Ly6a-independent AAVs (gray label background) packaging CAG-mNeonGreen and imaged 24 hours post transduction. An overlay of brightfield and fluorescence images is presented. Scale bars = 200 μm. **c,** Representative images of HEK cells cultured in 96-well plates, either mock-transfected (none) or transfected with Ly6a or Slco1c1. Slco1c1 infected cells display dim and diffuse fluorescence that extends beyond cell boundaries for every AAV tested. An overlay of brightfield and fluorescence images is presented. Scale bar = 200 μm.

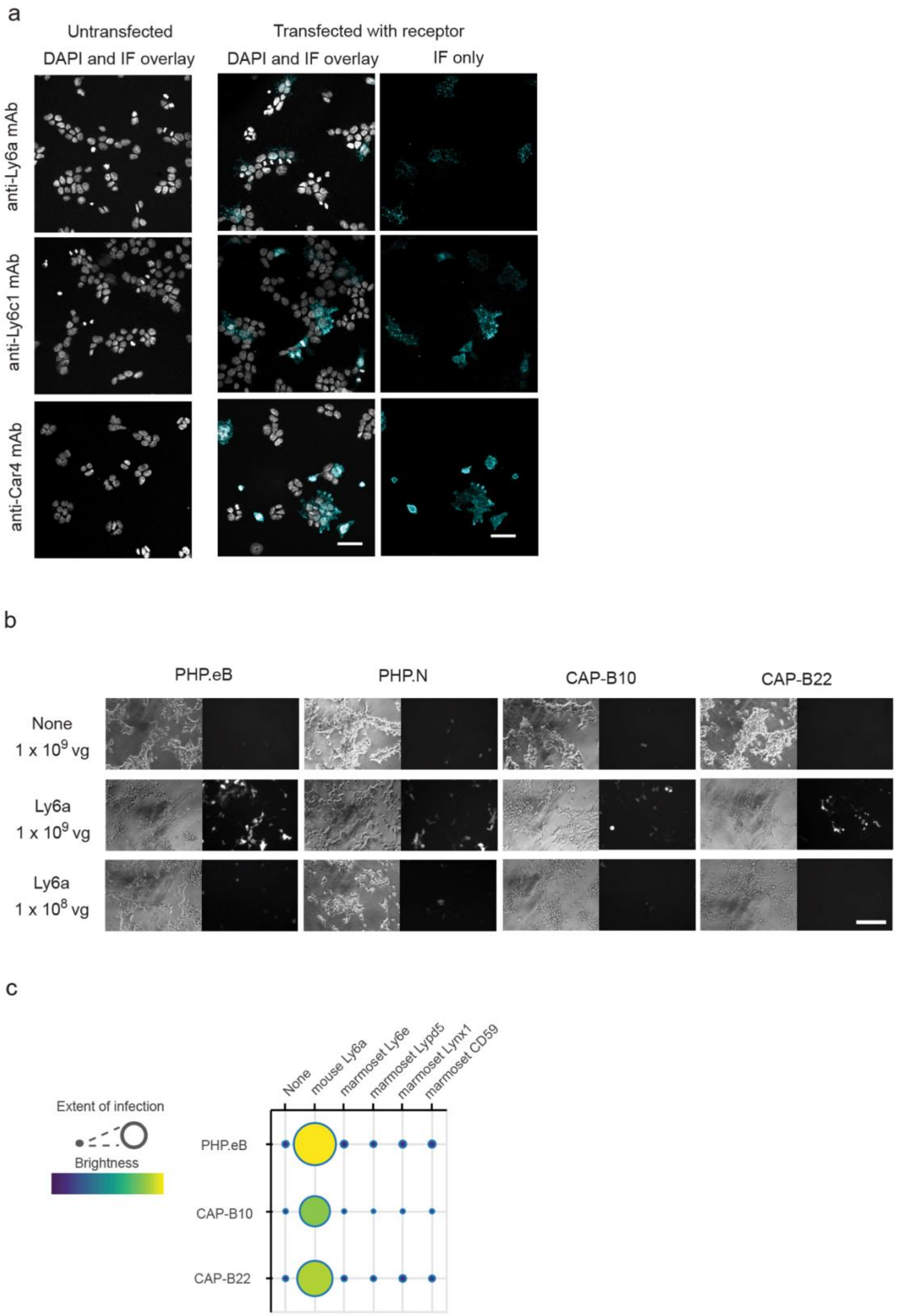


**Fig. S2a. Immunofluorescence confirms expression of selected membrane proteins from the**

**cell culture screen.** DAPI and immunofluorescence (IF) overlay for untransfected and receptor-transfected HEK293T cells stained with the indicated antibody as well as the IF channel alone from receptor-transfected HEK293T cell images (at left). Scale bars = 50 μm. **b. Ly6a-interacting AAVs engineered from PHP.eB display the same degree of potency enhancement in cell culture.** Brightfield (*left*) and fluorescence (*right*) images of PHP.eB, PHP.N, and CAP-B10 packaging CAG-mNeonGreen as well as CAP-B22 packaging CAG-NLS-GFP at two different doses in mock-transfected and Ly6a-expressing cells. Scale bar = 100 μm. **c. CAP-B10 and CAP-B22 do not display a potency enhancement with marmoset Ly6 proteins.** Potency of engineered AAVs for HEK293T cells transfected with the potential receptor panel. Extent of infection (Max: 0.52, Min: 0.05), Total brightness per signal area (Max: 0.79, Min: 0.17).

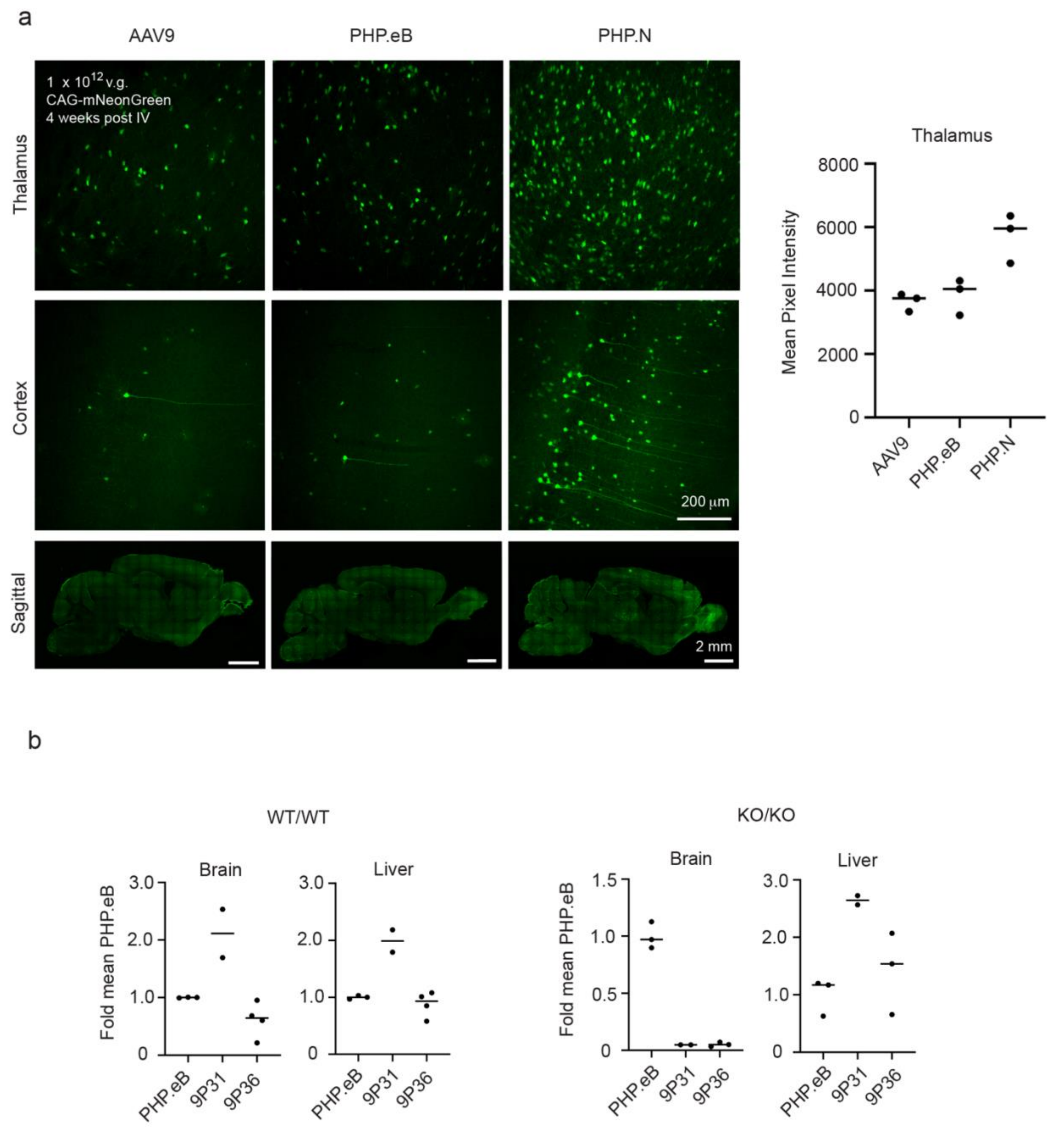


**Supplementary Figure 3a. In CBA/J mice at high doses, PHP.N outperforms AAV9 and PHP.eB.** Animals were retro-orbitally injected at 6-8 weeks old with 1 x $10^{12}$ v.g. of AAV9, PHP.eB, or PHP.N packaging CAG-mNeonGreen (n = 3). Four weeks after injection, tissue was collected and imaged to detect virus potency. Each data point in the thalamus mean pixel intensity quantitation represents one animal and bars indicate the mean value. **b. Quantification of AAV**

**potency in WT and Car4-KO mice.** Mean pixel intensity of whole brain sagittal slices and liver sections were quantified and each individual datapoint was then normalized to the mean PHP.eB intensity. Each data point represents one animal and bars indicate the mean value.

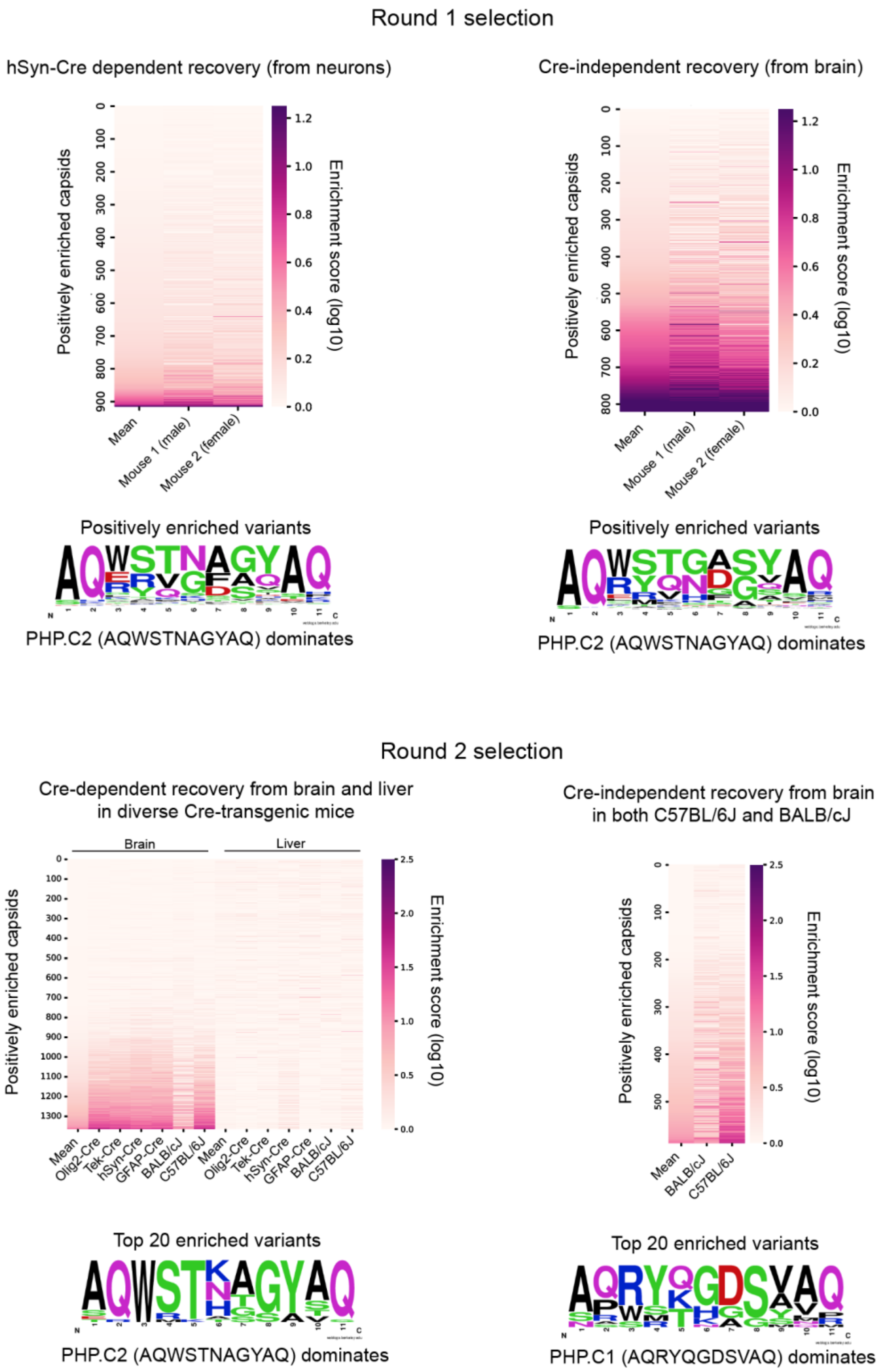


**Supplementary Figure 4. Enriched capsid variants during M-CREATE selections.** Capsids ranked according to their enrichment score from round 1 (top) and round 2 (bottom) selections using M-CREATE Cre-dependent recovery (left) or Cre-independent recovery (right). Sequence frequency plots indicate the patterns of the most enriched variants from each round and selection type.

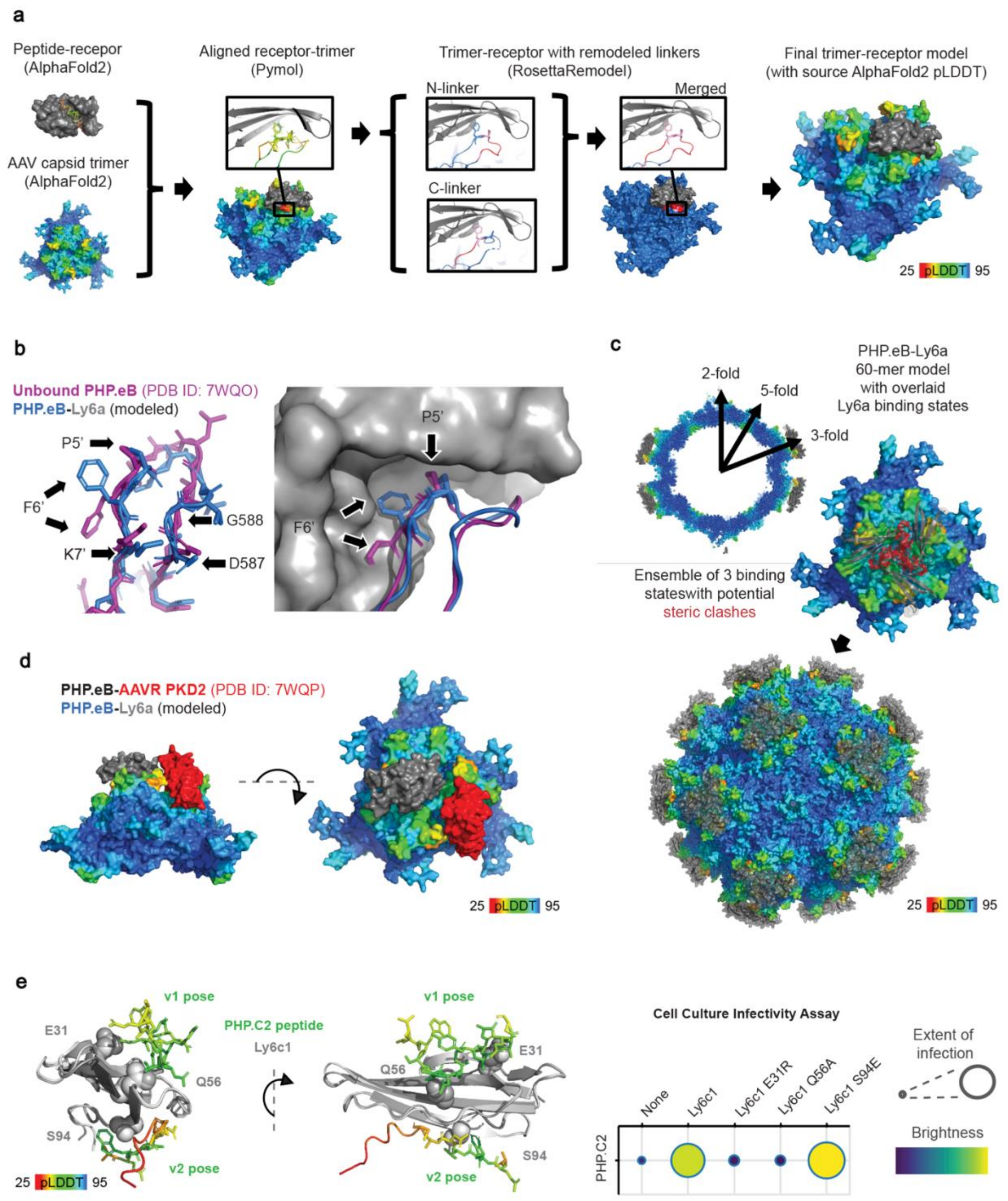


**Supplementary Figure 5. An integrative structure modeling method yields a snapshot of PHP.eB-Ly6a interaction. a,** A workflow for modeling engineered AAV-receptor complex structures (PHP.eB: colored by pLDDT score, Ly6a: grey). **b,** Comparison between computationally modeled PHP.eB-Ly6a (PHP.eB: blue, Ly6a: grey) and CryoEM structure of unbound PHP.eB (purple, PDB ID: 7WQO). All high-confidence residues (pLDDT > 45, see left panel arrow) within the inserted peptide showed consistent conformations between the two models except for F6’, which did not have clear side chain density in the CryoEM map. The side chain of F6’ predicted in the unbound PHP.eB model would cause a steric clash in the predicted complex model. **c,** Assembled PHP.eB capsid-Ly6a model representing an ensemble of all Ly6a binding states, despite steric clashes, as would occur in cryo-EM particle reconstruction (PHP.eB: rainbow, Ly6a:grey)**.** Top: cross section of the assembled capsid model. Middle: zoom-in of a 3-fold spike, highlighting steric clashes (red) between three different binding states. **d**, Overlaid structures of computationally modeled PHP.eB-Ly6a (PHP.eB: rainbow, Ly6a: grey) and CryoEM model of PHP.eB-AAVR PKD2 (AAVR PKD2: red, PDB ID: 7WQP). **e *left*,** Comparison between the computationally modeled PHP.C2-Ly6c1 binding poses predicted by AlphaFold-Multimer v1 and v2. PHP.C2 peptides are colored by residue according to pLDDT score. Ly6c1 residues chosen for mutation are shown as spheres. ***right,*** Potency of PHP.C2 for HEK293T cells transfected with wild type or mutant Ly6c1 receptors. Extent of infection (Max: 0.67, Min: 0.13), Total brightness per signal area (Max: 0.72, Min: 0.31).
